# Spatiotemporal transcriptome atlas of human embryos after gastrulation

**DOI:** 10.1101/2023.04.22.537909

**Authors:** Jiexue Pan, Yuejiao Li, Zhongliang Lin, Qing Lan, Huixi Chen, Man Zhai, Shengwei Sui, Gaochen Zhang, Yi Cheng, Yunhui Tang, Qingchen Wang, Ying Zhang, Fuhe Ma, Yue Xu, Yiting Mao, Qinfang Chen, Yichun Guan, Nan Meng, Haiqian Lu, Xiangjuan Li, Tingting Zheng, Xiaoying Yao, Qiuyu Qin, Bin Jiang, Yuxing Ren, Meiqi Luo, Ji Nancuo, Xin Jin, Jianzhong Sheng, Congjian Xu, Xinmei Liu, Yanting Wu, Chenming Xu, Lijian Zhao, Hongbo Yang, Ya Gao, Guolian Ding, Xun Xu, Hefeng Huang

## Abstract

The spatial and temporal atlas of gene expression in the human embryo at early gestation is critical in understanding embryo development, organogenesis, and disease origins. We obtained the spatiotemporal transcriptome from 90 sagittal sections of 16 whole human embryos from 3 to 8 post-conception weeks by Stereo-seq with high resolution and ultra-large field, establishing the development trajectory/regulatory profiling of 49 organs. We uncovered the organ-specific regulons as potential lineage-determining factors and identified the new regulatory networks during heart and brain development. The atlas refines the key organs/cell types vulnerable to virus infection and genetic disorders, and, reveals the dynamics of allelic gene expression in specific organs at different stages. These results present the first comprehensive delineation of the spatiotemporal transcriptomic dynamics of human organogenesis.

**One Sentence Summary:** The spatiotemporal transcriptome atlas presents a comprehensive delineation of human embryogenesis after gastrulation.

## Introduction

Human embryo Human embryo development begins with a fertilized ovum that divides and differentiates through preimplantation, embryonic, and fetal stages. After gastrulation, the embryo undergoes early organogenesis that intricately orchestrates in spatial dimensions and gestational age. Many childhood and adult disorders originate back from the embryonic stages, especially birth defects ^1^, metabolic disorders ^2^, and neurodevelopmental disorders ^3^. The periods most susceptible to teratogenesis are between 3 to 8 post-conception weeks (PCWs) when the embryo undergoes great expansion of cellular diversity and cell differentiation for organ morphogenesis ^4^. Thus, understanding the spatial and temporal expression of genes regulating embryogenesis and organogenesis is crucial in deciphering the etiology of pathological conditions, including congenital anomalies. Due to technical challenges and sampling difficulties, time-lapse studies of embryogenesis were mostly conducted in preimplantation embryos ^5–8^, and most of our knowledge of early human organogenesis was gleaned from conventional histological studies via embryo sectioning and reconstructions ^9, 10^.

Previously, great efforts were made to establish the gene-expression landscapes during early human embryogenesis. For instance, recent advancements in single-cell RNA sequencing (scRNA-seq) technologies have allowed a comprehensive analysis of human cellular heterogeneity in 15 fetal organs consisting of more than 4 million cells collected from 72 to 129 days of post-conception age ^11^, and, data from Xu et al. also provides a single-cell transcriptome atlas of early embryo ^12^. However, data crossing embryonic stages post-gastrulation remains unavailable. Furthermore, the analysis of intercellular regulatory networks and global transcriptional patterns in tissue spatial architecture was not performed. In recent years, spatial transcriptome technology was utilized to profile gene expression with spatial information in specific human fetal organs such as the digestive tract ^13, 14^, heart ^15^, liver ^16, 17^, lung ^18^, gonad ^19^, cortex ^20^, and cerebellum ^21^. These new data greatly advanced our understanding of cellular heterogeneity, complex tissue architectures, and dynamic changes of organogenesis during fetal development. It also underlines the significance of studying human samples, as model organisms cannot fully reflect the unique aspects of human development. Though two sagittal sections of a PCW5 human embryo were spatially mapped ^12^, there still lacks developmental dynamics of whole-embryo spatial transcriptome atlas due to excessive embryo size exceeding the limits of the most current spatial transcriptome technology. Hence, the molecular mechanisms of the cellular diversity and intricately orchestrated organogenesis in human embryo still requires better illustrations.

We have previously developed a spatial transcriptome method that allows the *in situ* profiling of gene expression with high signal resolution and ultra-large signal field. The method, known as Stereo-seq, exploits a modified chip containing the patterned DNA nanoballs (DNBs) randomly barcoded with molecular tags as the identity of spatial localization ^22^. Recently, Stereo-seq was used to build the spatial transcriptome maps in various model organisms, including the mouse embryos ^22^, *Arabidopsis* leaves ^23^, zebrafish embryos ^24^, *Drosophila* embryos ^25^, and axolotl whole brain ^26^. Here, we present a spatial and temporal transcriptome atlas of the whole human embryos from PCW3 to PCW8 at a 1-week temporal interval. The cellular heterogeneity and regulatory mechanisms underlying organ-specific specializations during human embryo development were explored. Spatial analysis in the heart, brain, skeletal muscle, liver, and spinal cord was conducted to identify regulatory networks during organogenesis. The atlas also provides evidence and refinement on existing knowledge of organ development and key organs/cell types that are vulnerable to virus infection and genetic disorders. Furthermore, we investigated the dynamics of tissue-specific allelic gene expression at different stages.

## Results

### Spatiotemporal transcriptomic atlas of human organogenesis

A total of 90 sagittal sections from 16 euploid human embryos (9 males, 7 females) were sequenced to obtain a spatial-temporal transcriptome atlas of organogenesis (Fig. 1A). The collected embryos ranged from the Carnegie stage (CS) 12 to 23, which were further categorized by PCW3-8 so that each PCW contained at least 2 embryos for further analysis (Table S1). Unsupervised spatially constrained-clustering (SCC) was performed with bin50 (50 × 50 DNB bins, equivalent to 25 μm in diameter), resulting in a total of 14,861,157 spots of all sections. An average of 2,819 unique molecular identifiers (UMIs) and 1,152 genes were obtained in each spot of bin50 (Table S1 and Fig. S1).

**Figure 1.**
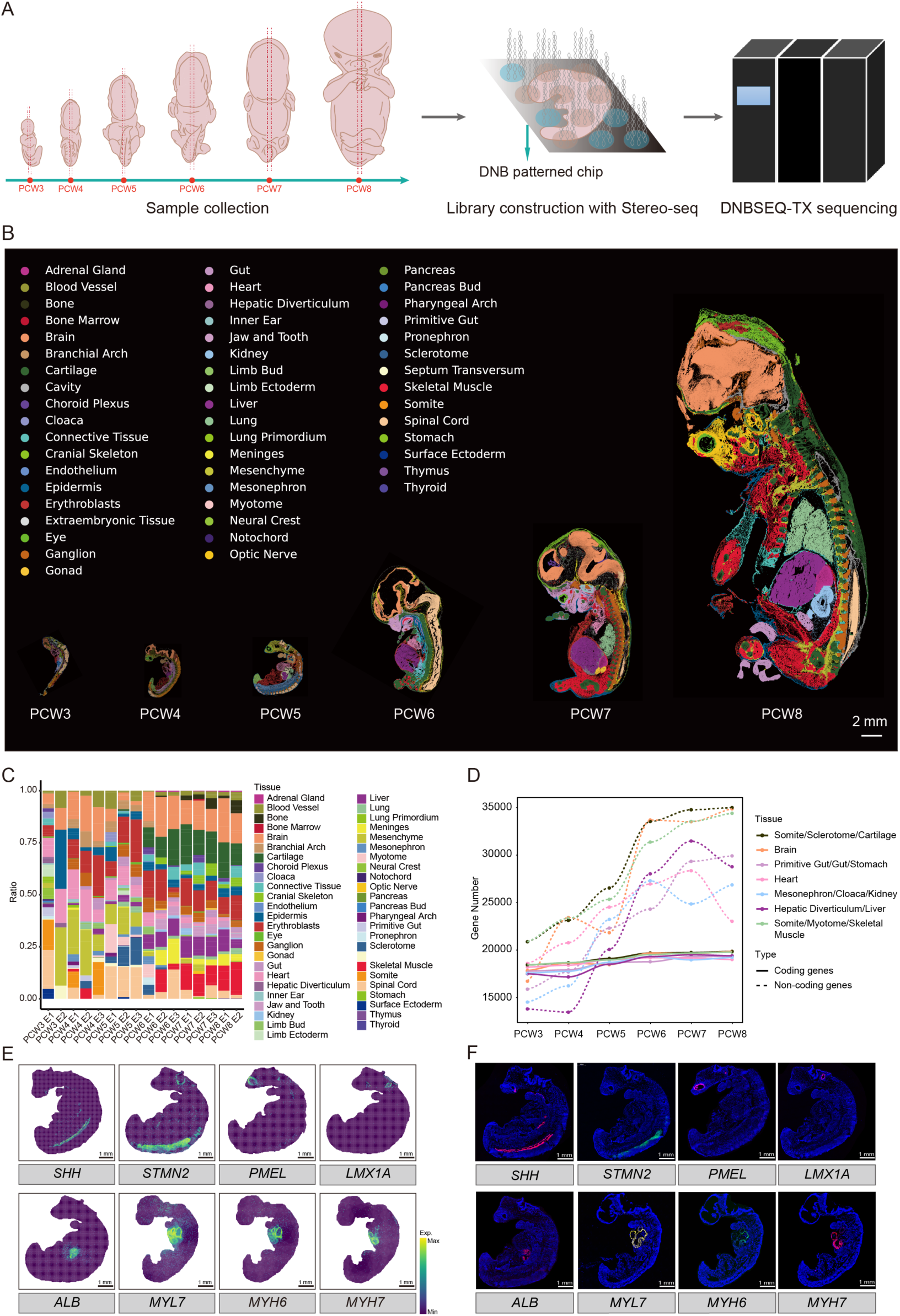
Spatiotemporal transcriptomic atlas of human organogenesis. (A) Schematic workflow of this study. Sagittal sections of human embryos across PCW3-8 were included for Stereo-seq. (B) Unsupervised SCC of human embryo sections across PCW3-8. Forty-nine anatomical tissues were annotated in indicated colors. (C) The percentage of bins annotated for each tissue shows the tissue-type distribution against samples. (D) The number of protein-coding and protein-non-coding genes in the main organs across 3-8 PCWs. (E-F) Spatial visualization of selected tissues using well-known tissue markers from PCW4 (*MYL7* in the heart, *MYH6* in the atrium and *MYH7* in the ventricle) and PCW5 embryos (*SHH* in the notochord, *STMN2* in the neural tube, *PMEL* in the eye, *LMX1A* in inner ear, *ALB* in the liver) on Stereo-seq maps (E) and corresponding RNA ISH images using adjacent sections (F). Scale bar, 1 mm.

As the embryo section size enlarged from 4 mm × 1.5 mm of PCW3 to 42 mm × 20 mm of PCW8, the number of sequenced reads and bin50 concordantly rose (Fig. S2A-B). Meanwhile, the number of uniquely mapped reads and annotated reads per bin50 remained relatively stable at different development stages (Fig. S2C-D), showing consistent data quality. Unsupervised spatially constrained-clustering (SCC) of bin50 Stereo-seq data was performed to identify diverse anatomic regions of the whole embryos. A total of 49 organs/anatomic regions were annotated using previously reported gene markers, including major organs (e.g. brain, eye, gut, heart, kidney, liver, lung, and spinal cord) and fine structures (e.g. inner ear, gonad, notochord, choroid plexus, etc.) (Fig. 1B). Plotting the tissue-type distribution against developmental stages showed that the heart, spinal cord, somite, and mesenchyme were the prominent tissue types in PCW3-5, whereas the liver, bone, cartilage, and skeletal muscle were prominent in PCW6-8 (Fig. 1C). During the transition from PCW3-5 to PCW6-8, the number of the protein-coding genes of multiple organs showed mild increases, while the number of non-coding genes surged sharply, particularly between PCW4-7 (Fig. 1D). To verify our unbiased clustering and annotation results, typical markers of multiple organs were stained by the *in situ* hybridization (ISH) experiments (*SHH* in notochord, *STMN2* in neural tube, *PMEL* in eye, *LMX1A* in inner ear, *ALB* in liver, *MYL7* in heart, *MYH6* in atrium and *MYH7* in ventricle), which showed good consistency to the Stereo-seq data (Fig. 1E-F).

### Spatiotemporal dynamics of representative Gene Regulatory Networks (GRNs) during human organogenesis

Using pySCENIC and Hotspot, over a hundred transcription factors (TFs) and regulon modules were identified in human embryos from PCW3-8 based on the activity scores and spatial coordinates (Fig. 2A and Fig. S3). With Stereo-seq data, we showed that the TFs and regulon modules expressed with specific spatial localizations, and that each main organ was consist of various numbers of regulon modules located at specific subregions (Fig. 2B-C and Fig. S3). As gestational age progressed, the regulon modules increased from 19 at PCW3 to 26 at PCW8 (Table S2). The organs with mostly enriched regulon expression changed from the spinal cord, brain, and heart at PCW3 (Fig. S3A-C) to the brain, cartilage, liver, and skeletal muscle at PCW8 (Fig. 2A-C). To facilitate the organ development analysis, we first determined the organ-identity regulators using the top organ-specific regulons with the highest regulon specificity score (RSS) of each organ (Fig. S4 and Table S3). This allowed the identification of classic regulators in the appropriate organs (CEBPA, SOX2, GATA4, and DRGX in the liver, brain, heart, and spinal cord, respectively) (Fig. S5C-F). Next, we compared organ identity regulators, and we found specific patterns of organ developmental trajectory from PCW3-8. For instance, MYF5 activity was observed in somite as early as PCW3, followed by the expression of a group of regulons including MYF5, MYF6, MYOD1, and MYOG in myotome at the tail and dorsal region at PCW5, and later enriched in skeletal muscle throughout the whole body at PCW7 (Fig. 2D and G). This suggests that the fate of skeletal muscle development is likely established around PCW3. Interestingly, we found that the alpha-actin gene *ACTA1* emerged at PCW7 in skeletal muscle when compared to the consistent expression of beta-actin gene *ACTB* and gamma-actin gene *ACTG1* throughout PCW3-8. In contrast, the High Mobility Group (HMG) genes including *HMGA1*, *HMGB1*, and *HMGB2* were strongly expressed in somite and myotome at PCW3-6 but silenced afterward (Fig. 2K). These distinct expression patterns suggest that different functional subunits and regulatory pathways are involved at different stages of fetal muscle development.

**Figure 2.**
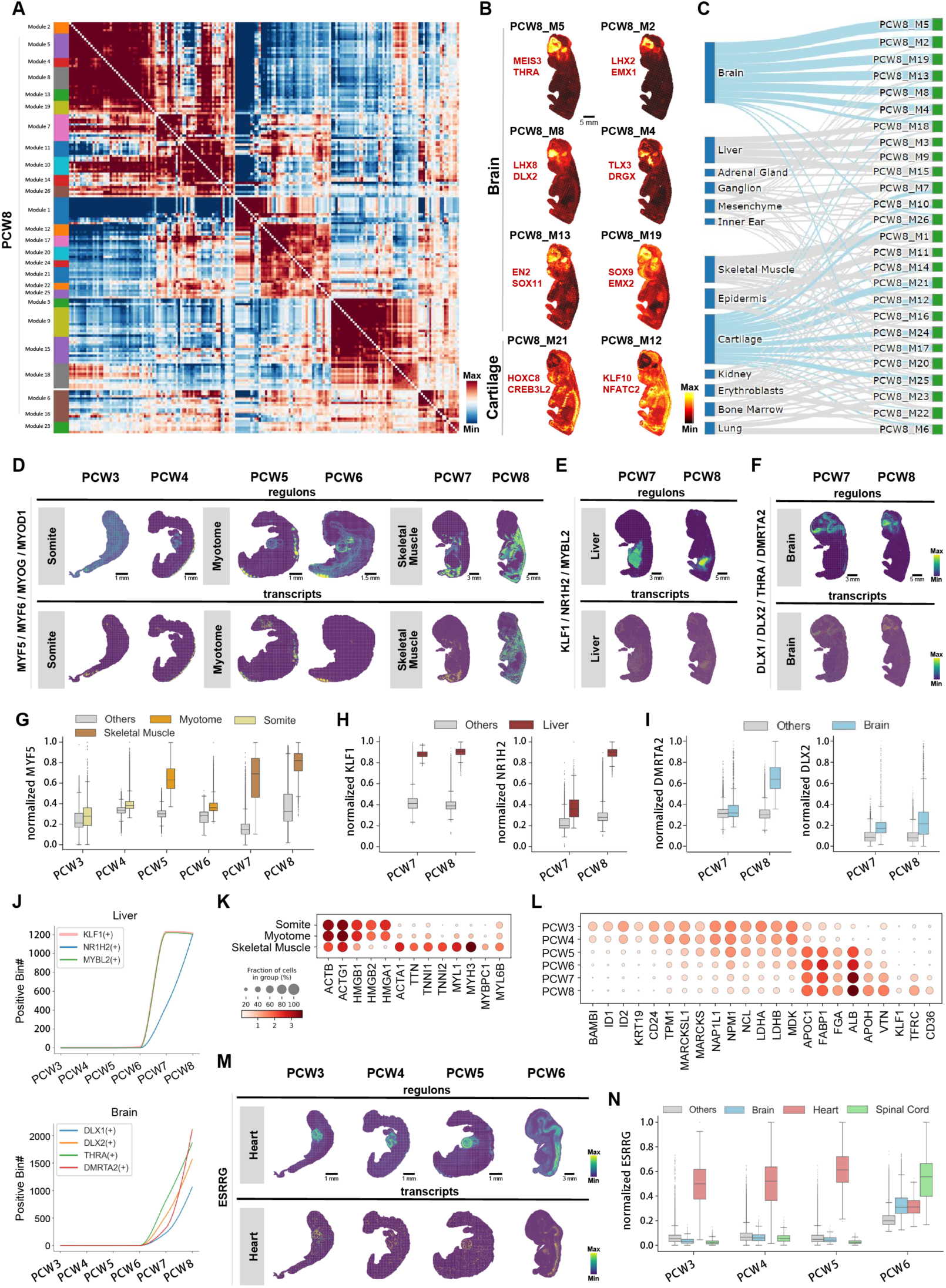
Comprehensive identification of GRNs in human organs across developmental stages. (A) Regulon modules grouped by Hotspot based on pairwise local correlation at PCW8. (B) Spatial patterns of representative regulon modules and two associated regulons in brain and cartilage at PCW8. (C) Sankey plot showing the spatial relationship between developmental organs and regulon modules at PCW8. Representative organs are highlighted in blue. (D) Spatial patterns of MYF5/MYF6/MYOG/MYOD1 regulon activities (top) and transcript expressions (bottom) from PCW3 to PCW8. (E) Spatial patterns of KLF1/NR1H2/MYBL2 regulon activities (top) and transcript expressions (bottom) from PCW7 to PCW8. (F) Spatial patterns of DLX1/DLX2/THRA/DMRTA2 regulon activities (top) and transcript expressions (bottom) from PCW7 to PCW8. (G) Boxplot showing MYF5 normalized regulon activity within somite/myotome/skeletal muscle and other organs from PCW3 to PCW8. (H) Boxplots showing KLF1 and NR1H2 normalized regulon activities within the liver and other organs from PCW7 to PCW8. (I) Boxplots showing DMRTA2 and DLX2 normalized regulon activities within brain and other organs from PCW7 to PCW8. (J) Line plot showing the total number of regulons KLF1, NR1H2, and MYBL2 bins in the liver from PCW3 to PCW8; line plot showing the total number of regulons DLX1, DLX2, THRA, and DMRTA2 bins in the brain from PCW3 to PCW8. (K) Bubbleplot showing the representative differentially expressed genes among somite, myotome, and skeletal muscle. (L) Bubbleplot showing the representative differentially expressed genes in the liver across developmental stages. (M) Spatial patterns of ESRRG regulon activities (top) and transcript expressions (bottom) from PCW3 to PCW6. (N) Boxplots showing ESRRG normalized regulon activities within the brain, heart, spinal cord, and other organs from PCW3 to PCW6.

In the liver, we also found an orchestrated regulation of differentially expressed regulons and genes. The regulon activities of TFDP1 (associating with hepatocytes proliferation and regeneration) ^27, 28^ increased after PCW7 and enriched in the liver (Fig. S5A), while the regulon modules of KLF1, NR1H2, and MYBL2 started to express at PCW7-8 in line with functions in erythroblast formation, cholesterol metabolism, and cell proliferation (Fig. 2E, 2H and Fig. S5A) ^29–31^. Meanwhile, several genes, including the lactate metabolic process genes of *LDHA*, *LDHB,* and erythroid progenitor developmental gene *ID1,* and *ID2* emerge and peak at PCW3-5 (Fig. 2L) ^32, 33^.

During fetal brain development, the regulons of DLX1, DLX2, THRA, and DMRTA2 surged since PCW6 (Fig. 2J) and were specifically activated at PCW7 (Fig. 2F, I-J and Fig. S5B), which is accordant with their roles in forebrain determination, neuronal and oligodendroglial determination (DLX1, DLX2) ^34^, brain development (THRA) ^35^, brain subregions specification, neuron and astrocyte determinations (DMRTA2) ^36, 37^. The orchestrated regulon expression and dramatic changes of brain structures indicate that PCW6-7 may be a critical stage for brain regionalization and function formation.

As one of the initial solid organs in human embryo, the heart displayed a remarkable level of the regulon activity of ESRRG, which is tied to the maturation of cardiac myocytes and maintenance of ventricular identity ^38, 39^, between PCW3-6 (Fig. 2M-N). It’s noteworthy that this regulon was further activated in tissues of brain and spinal cord at PCW6, which plays a crucial role in the oxidative glycolytic metabolism of neurons ^40^. The spatiotemporal dynamic changes in ESRRG highlight its pivotal roles in the regulation of early heart ventricular determination and, later on, in the metabolic capacity of central nervous system neurons.

### Cardiac development revealed by GRNs at the substructure level

To gain further insight into the regulatory networks governing the development of the heart chambers, we utilized the classic atrium (*MYH6, MYL7 and NR2F2*) and ventricle (*MYH7, MYL2 and SHPB7*) markers to annotate the cardiac substructures in multiple sections of PCW3-8 (Fig. 3A and Fig. S6A). We then compared the co-expressed genes in the atrium and ventricle, and identified top genes with distinct expressions at different stages (Fig. 3B and Table S4). Among these genes, the spatial expression patterns of genes highly expressed in the atrium (*COL2A1* and *ACTA1*) and ventricle (*SLC8A1* and *SORBS2*) were exhibited in Fig. S6A. Using pySCENIC, we identified several atrium-/ventricle-specific regulators, including the well-known ventricle-specific TFs TBX20 and ESRRG (Fig. 3C). Regulon ETS1 demonstrated high consistency with the trabecular cardiomyocytes’ spatial expression pattern (Fig. S6D).

**Figure 3.**
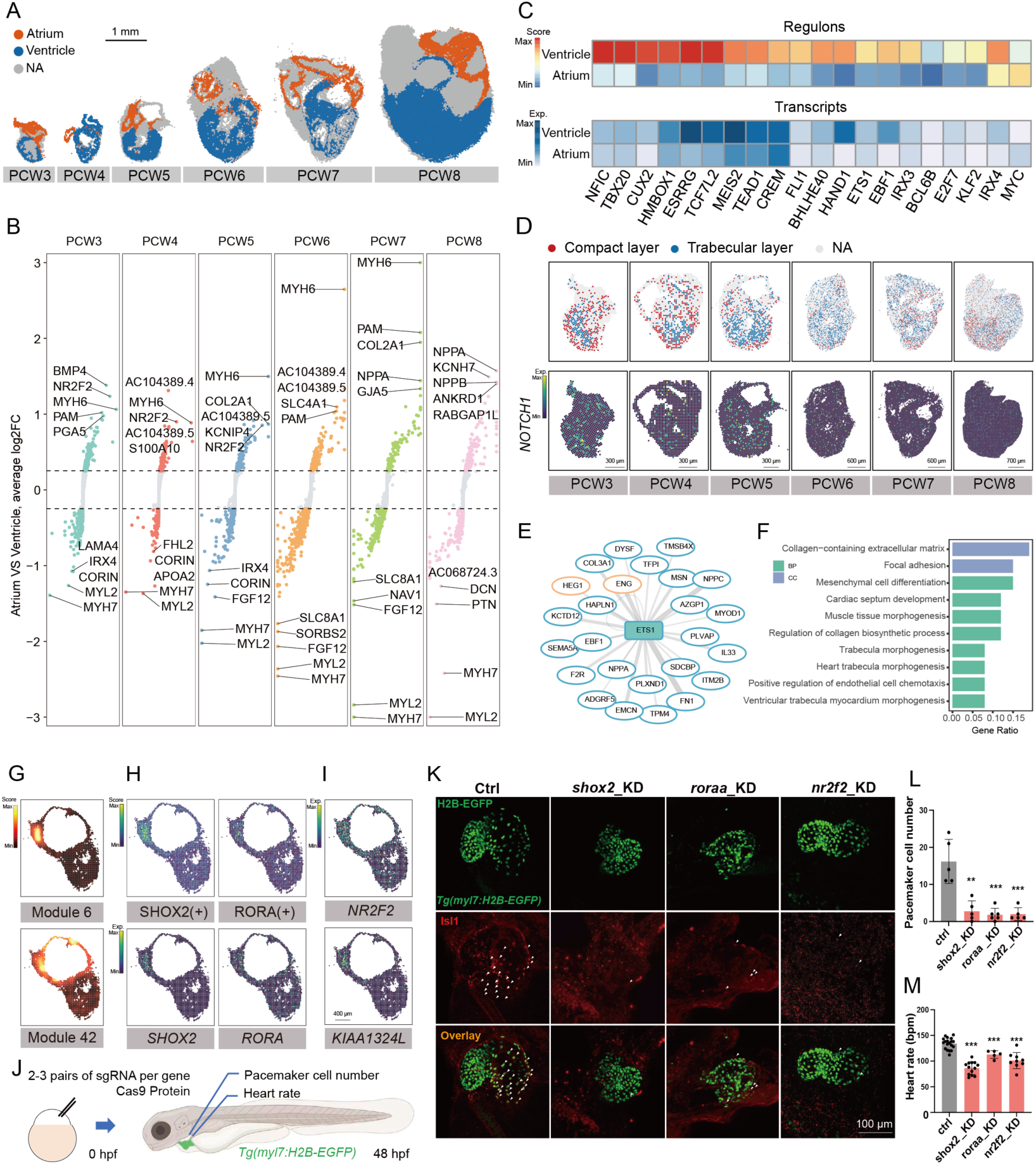
Cardiac development revealed by GRNs at the substructure level. (A) Unsupervised SCC and annotation of heart sections across PCW3-8. Scale bar, 1 mm. (B) Differentially expressed genes (DEGs) identified in the atrium and ventricle from all chips carrying heart tissues across stages. (C) Heatmap showing the enrichment of atrium-/ventricle-specific regulons identified by pySCENIC in the heart and their corresponding expressions. (D) The annotation of the compact layer and trabecular layer (top) and the spatial expression of NOTCH1 (bottom) in the heart across PCW3-8. (E) The regulatory network of ETS1, in which genes highlighted with orange circles represent genes for trabecular morphogenesis. The line width represents the weight of motif enrichment. (F) Barplot exhibiting the representative Gene Ontology enrichment terms of genes in E. (G) Spatial visualization of gene modules related to SAN in PCW6 heart. (H) The spatial visualization of regulon activity and gene expression of SHOX2 and RORA in PCW6 heart. (I) The spatial expression of *NR2F2* and *KIAA1324L* in PCW6 heart. (J) Schematic diagram of knockdown experiment in zebrafish. (K) Immunohistochemistry of SAN in zebrafish at 48 hpf. Pacemaker cell nuclei are Islet-1 (Isl1) + (red) in *Tg(myl7:H2B-EGFP)* transgenic fish (denoted by white arrows). (L) The pacemaker cell number in control, *shox2*, *roraa,* and *nr2f2* knockdown zebrafish at 48 hpf. **p < 0.01, ***p < 0.001. (M) The heart rate of control, *shox2*, *roraa,* and *nr2f2* knockdown zebrafish at 48 hpf. ***p < 0.001.

Trabeculation is a critical morphological milestone in the development of ventricular chambers, yet the regulatory processes are only partially understood. To shed light on this, we employed a set of genes associated with cellular differentiation and tissue remodeling during trabeculation (as detailed in the methods) as a trabecula gene module. We then used this module to determine the localization of the trabecula layer in the heart, which exhibited temporal and spatial consistency with *NOTCH1*, one of the most thoroughly investigated signaling axes in cardiac trabeculation ^41^ (Fig. 3D). Interestingly, this trabecula gene module was activated during PCW3-5, pointing to a possible crucial time window of trabecular formation (Fig. S6C). Consistently, the regulon activity of ETS1 was activated between PCW3-5 at the trabecular layer and co-localize with *NOTCH1* at PCW3-5 (Fig. S6D). By identifying ETS1 target genes (Fig. 3E), we found that *ENG* and *HEG1* were involved in heart trabecula morphogenesis (Fig. 3F), thus providing further evidence of the important role of *ETS1* in regulating cardiac trabeculation.

The sinoatrial node (SAN) is a small and specialized structure situated at the junction of the superior vena cava and right atrium. The high-resolution of Stereo-seq enabled us to pinpoint the location of the SAN in the embryonic heart, providing a better understanding of GRNs in human SAN development. In PCW6, we identified two modules that exclusively localized at the (SAN using Hotspot analysis (Fig. 3G and Table S5). These modules contained *SHOX2*, a key regulator of pacemaker differentiation ^42^, and *VSNL1*, a core GRN governing the function of SAN in mice ^43^. PySCENIC’s GRNs analysis further demonstrated the specific regulons of SHOX2 and RORA in the SAN (Fig. 3H). RORA has been reported as a pivotal TF in regulating circadian rhythms ^44^. The downstream genes of SHOX2 (*SLIT2*, *ATP2A2*, *TBX18*, *PPP1R1A*, *TBX5*, *FAM78A*, *CIRBP*, *PRKG1)* and RORA (*PRKG1*, *ZNF385B*, *MAST4*, *DGKI*, *PGM5P4-AS1*, *RERE*) were also spatially expressed in the SAN region of the PCW6 heart (Fig. S7A-B). Meanwhile, the spatial gene modules further identified *NR2F2* and *KIAA1324L* as SAN-specific genes (Fig. 3I, Fig. S7C). To verify their regulatory roles in SAN development, the target gene was knocked down in *Tg(myl7:H2B-EGFP)* transgenic fish with specifically labeled cardiomyocyte nuclei ^45^ (Fig. 3J). Knockdown of *SHOX2*, *RORA*, *NR2F2*, *KIAA1324L* and *VSNL1* orthologs in zebrafish resulted in a significant reduction of pacemaker cell number labeled by Islet-1 (Isl1) in 2 days post fertilization (dpf) embryos (Fig. 3K-L, Fig. S8A-B and movie S1-7). Moreover, the knockdown of these genes in zebrafish embryos all reduced heart rate (Fig. 3M, Fig. S8C and movie S8-14). These data collectively support the regulatory roles of *RORA, NR2F2*, *KIAA1324L* and *VSNL1* in SAN development.

### Cellular and molecular changes in the regionalization of the human embryonic nervous system

The development of human nervous system starts from the embryonic stage and extending postnatally throughout infancy, childhood, adolescence, and young adulthood. Myriads of functionally diverse cell types, circuits, and regions are formed over time. A dataset from BrainVar, involving 176 human frontal cerebral wall samples across prenatal and postnatal development, indicated that early brain development coincides with the establishment of regional identity across the brain ^46^. However, the cellular and molecular spatial characterizations of fetal brain development at the embryonic stage are rare. Using previously reported markers ^47–50^, brain subregions such as forebrain (Fb), midbrain (Mb), hindbrain (Hb), and spinal cord (SpC) were spatially projected. Additionally, pallium ventricular zone (Pall VZ), pallium (Pall), subpallium ventricular zone (SPall VZ), subpallium (SPall), optic recess (Or), diencephalon (Die), and hypothalamus (Hy) in Fb, Mb ventricular zone (Mb VZ) in Mb, Hb ventricular zone (Hb VZ) and cerebellum (Cere) in Hb were spatially designated from PCW3 to PCW8 (Fig. 4A, B and D). The number of the annotated brain regions and subregions rapidly grew from 5 to 12 through PCW3-8, and the proportions of several subregions including Spall, Pall, Mb, and Cere in the whole brain dramatically increase after PCW6 (Fig. 4B). Thus, PCW6 may be a key period of fetal brain diversification and specialization. This is further supported by the correlation analysis of the transcriptional similarity among different brain regions from PCW3 to PCW8 (Fig. 4C). Overall, the brain regions at the identical gestational stages shared the strongest correlations. The clustering peaked between PCW3-5 and weakened afterward, indicating the heterogeneous differentiation and increased functional complexity of brain subregions occurred from PCW6.

**Figure 4.**
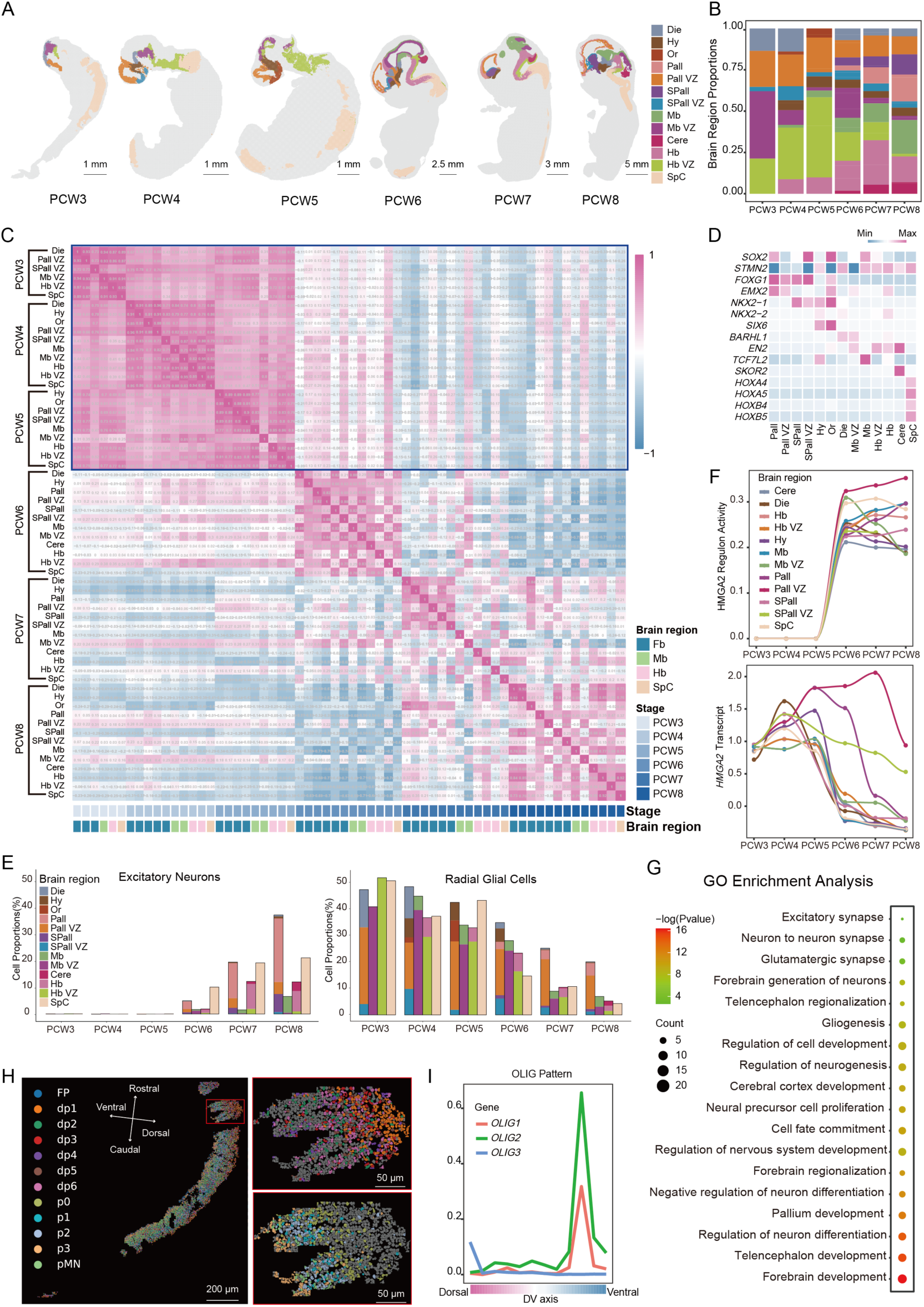
Cellular and molecular changes in the regionalization of the human embryonic nervous system. (A) Spatiotemporal transcriptomic atlas of the human embryonic nervous system. (B) The percentage of bins annotated for brain regions. (C) The heatmap shows the Spearman correlation between brain regions at each stage. (D) Heatmap shows the marker gene expression in regions of the human embryonic nervous system. (E) The bar plot shows the cell proportion of excitatory neurons and radial glial cells in each brain region (the height of the bars indicates cell proportion in the Fb, Mb, Hb, SpC, and the color of the bars indicates the composition ratio of fine brain regions). (F) Temporal dynamics of HMGA2 regulon activity and expression. (G) GO enrichment analysis of HMGA2 target genes. (H) Annotation of anatomical structures in PCW3 spinal cord at single-cell resolution. (I) The dorsoventral axis pattern of the OLIG family in the spinal cord.

The percentage of each cell type was calculated according to the marker gene (Table S6) to further characterize the landscape of fetal brain development. Both excitatory and inhibitory neurons dramatically emerged at PCW6-8, which is consistent with the progression of neurogenesis within this time. Nonetheless, a small number of inhibitory neuron markers can be detected as early as PCW3, while the marker of excitatory neurons emerges at PCW6 and afterward (Fig. 4E and Fig. S9D), suggesting delayed neurogenesis of excitatory neurons in comparison to the inhibitory neurons. The excitatory neurons are mainly located in Pall in the forebrain, while inhibitory neurons in the ventral ganglion eminence (e.g. SPall, Hy, Die, etc.) (Fig. S10F, H-I), which is accordant with previous studies ^51^. In progenitor cells, the proportions of radial glial cells, neural progenitor cells, and oligodendrocyte progenitor cells all showed a declining tendency over time (Fig. 4E, Fig. S9A-B and Fig. S10B-E), whereas the glial progenitor cells underwent a steady proliferation during PCW3-8 (Fig. S9C and S10G). The spatial annotation of different progenitor cells was consistent with their localization in specific subregions (Fig. S10B-E and G). However, the development of glial cells was relatively invariant during PCW3-8 (Fig. S9E-G and S10J-L).

Among the TFs and regulon modules during embryonic nervous system development, we identified 20 regulons specifically enriched in PCW6-8 but not in earlier stages (Fig. S11A-B). Specifically, *HMGA2* was ubiquitously expressed in the whole brain with no regulon activity during PCW3-5 but then specifically concentrated in Pall VZ with regulon activity at PCW6-8 (Fig. 4F and Fig. S11C-D). The downstream target genes of HMGA2 were mainly related to forebrain development, forebrain regionalization, gliogenesis, and forebrain generation of neurons, glutamatergic synapse, and excitatory synapse in PCW6-8 (Fig. 4G, Fig. S11E and Table S7). Therefore, HMGA2 may be identified as a key regulator of cerebral cortex development.

Previously, a dorsal-ventral expression pattern was reported in the fetal spinal cord ^52^. We segmented the cell boundaries with *spateo* and aligned the spinal cord Stereo-seq data of PCW3 with previous scRNA-seq data ^52^ (Fig. 4H). We found similar expression patterns of marker genes used to identify dorsoventral (DV) domains in human progenitors (Fig. S9H). Additionally, we found that OLIG1 and OLIG2 were mainly located at the ventral side of the spinal cord, while OLIG3 was at the dorsal side (Fig. 4I, Fig. S9L-N and Fig. S10D-E and M). Furthermore, we showed that the spatial localization of specific cell types in the spinal cord also displayed the dorsal-ventral distribution. Radial glial cells were located on both sides of the dorsal-ventral axis of the spinal cord (dp1, FP, and p3), with neural progenitor cells in the center (dp6, p0) and oligodendrocyte progenitor cells in pMN (Fig. S9I-L and S10B-E).

### Organ and cell type vulnerability to viruses and developmental disorders

Early pregnancy is a period with increased susceptibility to pathogens and congenital diseases. To improve our understanding of the interactions between pathogens and fetal host during early pregnancy, we identified the enriched expressions of receptors for 14 viruses (Table S8) with tissue-and time-specific patterns (Fig. 5A and Fig. S12A). For instance, *NECTIN1* was enriched in the eyes during PCW3-5 (Fig. 5A), which is consistent with the gene’s function to mediate the entry of the Herpes simplex virus (HSV) into the cornea ^53^. In the fetal liver, *NTCP*, the known receptor of the hepatitis B virus (HBV) ^54^, maintained a low expression level in the liver and other organs at PCW3-8 (Fig. 5A and Fig. S12B), while the receptors of hepatitis C virus (HCV), including *ASGR1* and *CLDN1*, are enriched in liver at PCW3-8 (Fig. 5A-B). Interestingly, several potential HBV receptors, including *ASGR1* ^55^ and *TFRC* ^56^, exhibit liver-specific expression from PCW3 and PCW6, respectively (Fig. 5A-B), supporting the possibility of mediating HBV infection via alternative pathways. Furthermore, we found the absent expression of the severe acute respiratory syndrome coronavirus 2 (SARS-CoV-2) receptors ^57^ in the fetal lung across PCW3-8 (Fig. 5A and Fig. S12B). Instead, *TMPRSS2* and *ACE2* were excessively co-expressed in the fetal gut at PCW8 (Fig. 5A). Main cell types in the gut, including epithelia, endothelia, neurons, mesenchyme, myeloid, red blood cells, B cells, and T cells, were annotated by both cell-type deconvolution ^58^ and cell markers identification ^13^ (Fig. 5C and Fig. S12C). Both the spatial localization (Fig.5C-D and Fig. S12D) and cell type contribution (Fig. 5E) showed that *ACE2* and *TMPRSS2* were enriched in gut epithelia at PCW8, but rarely co-expressed in the same cell (Fig. S12E).

**Figure 5.**
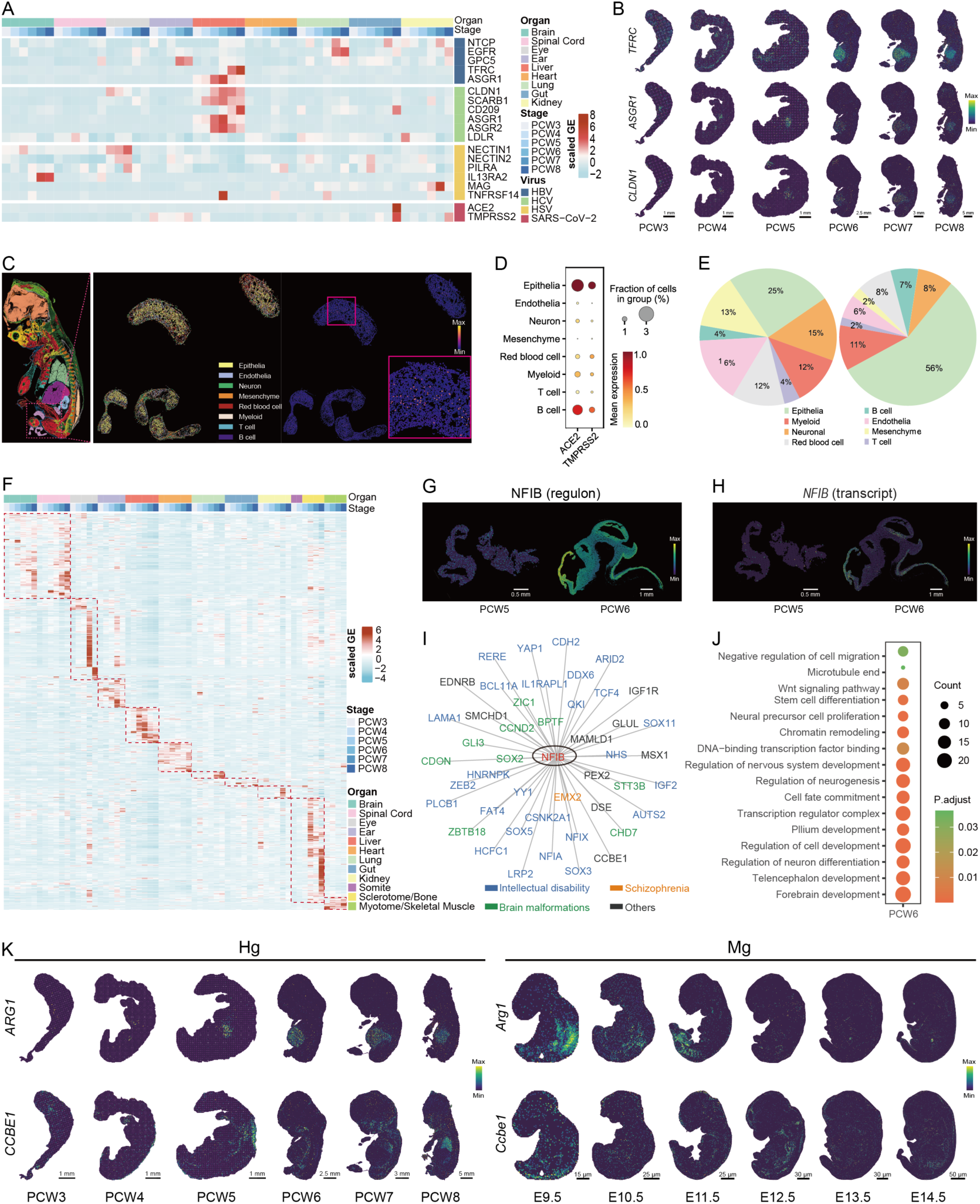
Association with virus infection and human developmental disorders. (A) Heatmap showing the normalized expression level of receptors for HBV, HCV, HSV, and SARS-CoV2 in 9 representative anatomic regions (brain, spinal cord, eye, ear, liver, heart, lung, gut, kidney) in embryo sections from PCW3 to PCW8. Embryo sections including PCW3 E1S3, PCW3 E1S5, PCW3 E1S8, PCW4 E1S4, PCW4 E1S7, PCW4 E3S4, PCW5 E3S1, PCW5 E3S2, PCW5 E3S3, PCW6 E1S10, PCW6 E2S1, PCW6 E2S2, PCW7 E2S4, PCW7 E2S5, PCW7 E1S4, PCW8 E2S5, PCW8 E2S6 and PCW8 E2S9 were used in this analysis. (B) Spatial visualization of the expression patterns of TFRC/ASGR1/CLDN1 from PCW3 to PCW8. (C) Spatial visualization of cell types and ACE2 expression based on single-cell segmentation in the gut at PCW8. (D) Bubbleplot showing the normalized expression of ACE2 and TMPRSS2 expression in the indicated cell types. (E) The proportion of cell types of all gut cells versus cells with ACE2 expression. (F) Heatmap shows the normalized expression level of 1,225 of the 1,922 genes selected from the developmental disorders genotype-to-phenotype database (DDG2P) in the representative anatomic regions (brain, spinal cord, eye, ear, liver, heart, lung, gut, kidney, somite, sclerotome/bone, and myotome/skeletal muscle) in embryo sections from PCW3 to PCW8. Embryo sections are the same as those in Fig. 5A. (G) Spatial visualization of NFIB regulon activity in the brain at PCW5 and PCW6. (H) Spatial visualization of NFIB transcript expression in the brain at PCW5 and PCW6. (I) Gene regulatory networks of NFIB in the brain at PCW6 as visualized by Cytoscape. Selected target genes in the DDG2P database were shown. (J) Bubbleplot showing the GO enrichment pathways of 171 target genes of NFIB in the brain at PCW6. (K) Spatial visualization of ARG1 and CCBE1 expression in the human embryo from PCW3 to PCW8, Arg1 and Ccbe1 expression in mouse embryo from E9.5 to E14.5.

To demonstrate the critical pathogenic windows of organs susceptible to genetic variations, we explored the expression landscape of 1922 genes associated with 1735 developmental diseases (Table S9) from the genotype-to-phenotype database (DDG2P V3.0) ^59^. The majority of these genes were mainly enriched in certain tissue at specific developmental stages, including the brain, spinal cord, eye, ear, liver, heart, lung, bone, and muscle (Fig. 5F and Table S10). Additionally, we identified time-specific gene expression in small subregions. For example, a TF gene *NFIB* expressed in several fetal organs such as brain and lung across PCW3-8 (Fig.S13A), and after digging into the subregions of brain, we found the expression exhibited spatiotemporal specificity, displaying a boost from PCW5 to PCW6 in Pall and Pall VZ of fetal brain, and accordant to its regulon activity (Fig. 5G-H and Fig. S13A-B).

Meanwhile, the number of NFIB downstream genes also dramatically increased within this time window (Fig. 5I, Fig. S13C and Table S11-12), which involves the pathways of the pallium and forebrain development, microtubule end, chromatin remodeling, and so on (Fig. 5J and Table S13). These results are consistent with the current evidence of NFIB in developing intellectual disability and macrocephaly ^60^.

To demonstrate the uniqueness of our data from animal models in genetic disease research, we compared the expression pattern of 1922 developmental diseases-associated genes between mouse embryos ^22^ and human embryos. We found that most results are consistent although some discrepancies were identified (Table S14). For instance, *ARG1*, a mutation causing argininemia or arginase-1 deficiency, and growth retardation and intellectual disability ^61, 62^, was specifically expressed in the fetal liver from PCW5 (Fig. 5K). Meanwhile, up-regulated genes in the human liver associating with arginine-1 deficiency pathways could be identified (Fig. S13D and Table S15). In mice, on contrary, the ortholog of *ARG1* in the liver or other organs from E9.5-E14.5 (Fig. 5K) displayed no obvious enrichment. In another example, *CCBE1*, known to regulate lymphatic vascular development ^63^, showed strong lung-specific expression in human at PCW8. In mice, *CCBE1* ortholog is minimally expressed in the lung from E9.5-E14.5 (Fig. 5K). Consistently, among the up-regulated genes in the human lungs compared with mice, pathways for lymphatic development were enriched (Fig. S13E and Table S15). Overall, these results illustrate the expression differences of disease-associated genes across species and emphasize the caution of applying mouse models for certain human disease studies.

### The dynamics of allelic gene expression in early embryo development

In most circumstances, both alleles of a gene are transcribed, although some genes show monoallelic expression. Selected expression of the two alleles determined by the parental origin is known as imprinting, and its dysregulation is often involved in growth disorders and neurological disorders ^64^. The expression pattern of the imprinted genes in early human embryo development remains largely unknown, and we systematically investigated the expression pattern of the monoallelic genes from PCW4 to PCW7 with spatial transcriptome data of fetal embryos. Using the spatial transcriptome of 15 embryo sections from 4 embryo samples (Fig. 6A and Table S16), a total of 555 genes were initially determined as tissue-specific monoallelic expression genes (Table S17). Next, the whole genomic sequencing of PCW4 Embryo3 and PCW7 Embryo3 with corresponding maternal decidua and the third-generation sequencing of the embryo samples were combined to establish embryo haplotypes. The embryo haplotyping results were then compared to the tissue-specific transcriptome data to identify the parental origins of allelic expression (Fig. 6A). After filtering low-quality results, a total of 114 monoallelic expression genes were identified, including 16 imprinted genes and 98 genes with parental origins in specific organs (Fig. S14 and Table S18). The allelic expression pattern of imprinted genes is verified as expected, such as *IGF2* and *MEST* with paternal allelic expression, and *MEG3* with maternal allelic expression (Fig. 6B). Meanwhile, the monoallelic expression pattern of *MEST* and *MEG3* was spatially consistent with their transcript localization (Fig. 6B), which validates unbiased analysis of monoallelic expression against the transcript expression analysis. From PCW4 and PCW7 embryos, we also found 6 genes showing parent-of-origin differential expression in certain tissues across multiple embryos (Fig. 6C-F, Fig. S14 and Table S19). Cytoplasmic poly(A)-binding protein 1 (*PABPC1*), an important modulator in mRNA post-transcriptional regulation, shows maternal-allelic expression across all tissues in embryos of different stages (Fig. 6D). *RN7SKP255*, the pseudogene of *RN7SK*, also shows high maternal-allelic expression (Fig. 6E). We also found that *GFPT2* and *SCIRT* show stronger paternal-allelic and maternal-allelic expression in the brain, respectively (Fig. 6F).

**Figure 6.**
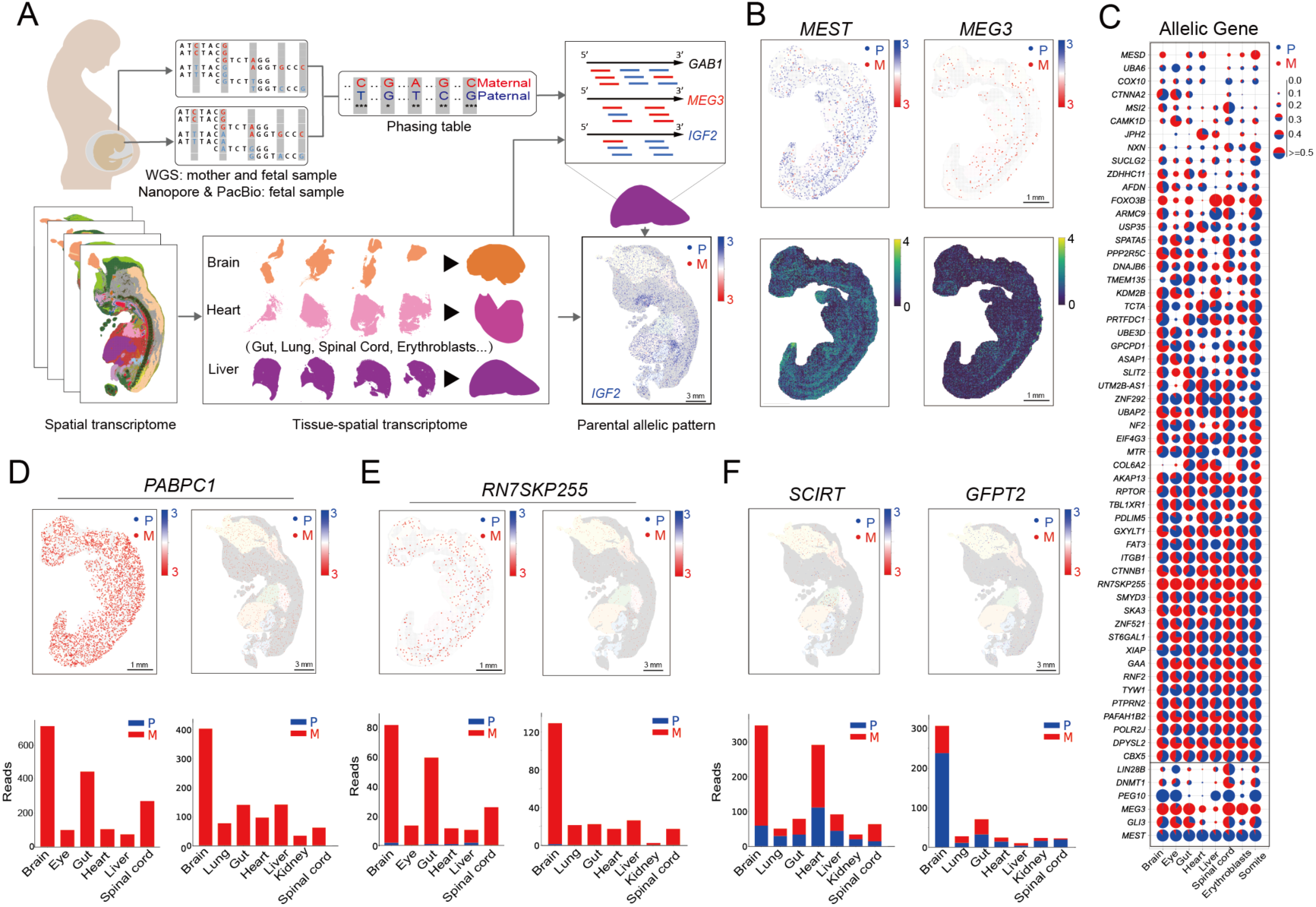
The existence of parental special expression pattern. (A) The allelic expression analysis flowchart of spatial transcriptome from multiple chips. (B) The spatial allelic expression pattern (upper) and common expression pattern (lower) of MEST and MEG3. (C) The parent-of-origin expressions of 60 phased genes in 8 tissues of PCW4 embryo. (D-E) The spatial maternal-allelic expression pattern of PABPC1 and RN7SKP255 in PCW4, accompanied by the validation of PCW7. (F) Brain-specific paternal-allelic and maternal-allelic expression pattern of GFPT2 and SCIRT in PCW7 embryo.

## Discussion

The Human Developmental Cell Atlas (HDCA) initiative aims to build a comprehensive atlas of human development at cellular resolution ^65^ and has a profound impact on biology and medicine by bringing a better understanding of anatomy, physiology, pathology, and intra/inter-cellular regulation in developmental human beings. The ethical 14-day rule, which limits the culture and study of intact human embryos to 14 days post-fertilization, keeps the embryogenesis after gastrulation still a black box ^66^. The human embryo undergoes early organogenesis after gastrulation, which requires precise spatiotemporal transcriptional regulation. During this period, most named parts of the body become identifiable, and many congenital birth defects and developmental disorders may also originate from this susceptible window of development. In recent years, powerful approaches have emerged which enabled the charting of dynamic changes during human development at single-cell resolution ^11, 12, 17, 67–72^, and recent spatial transcriptome data illustrated spatial transcriptional characteristics and complex architectures of certain developing organs ^14, 15, 18, 19^. Xu et al. also provided a spatially mapped view of a PCW5 embryo ^12^. In this study, our characterization provides a panoramic transcriptome atlas of *in vivo* human embryogenesis after gastrulation in both spatial and temporal resolution. This is the first time-lapse transcriptomic map to date portraying the molecular dynamics of organogenesis in the background of spatial localization in a complex, developing human embryo. *In situ,* gene expression of tissue sections as large as 15 cm^2^ can be realized owing to the large vision field and ultra-high density of the DNB probe. Gene expression profiles combine spatial information to help define developing systems. As illustrated in the atlas, great expansions of organ diversity in a whole embryo and cellular heterogeneity in the main organ occur during the transition from PCW3-5 to PCW7-8.

The susceptibility window of embryos to teratogens is usually an obscure period obtained by epidemiological statistics, which lacks molecular evidence support ^66^. Our spatial and temporal atlas will provide evidence and refinement of existing knowledge on embryonic development, as well as facilitate the delineation of fine structures and the descriptions of developmental trajectories of various systems. Therefore, this work helps to interpret the cellular and molecular mechanistic basis of developmental flaws in genetic disorders and nongenetic congenital diseases. Identification of homologous gene transcription in developing human embryos will allow discoveries in model organisms to be bridged to humans.

Genetic regulation of embryogenesis involves a cascade of genes and a nested pattern of transcriptional factors to create various cell types. In order to define whether the regulatory mechanisms during organogenesis was due to differences in cell type composition or transcriptomic differences, we identified the expression level of key transcriptional factor and comprehensively mapped GRNs. As expected, each organ is associated with multiple regulon modules and manifested a dynamic change, and the number of identified regulon modules also increases as gestational age progresses. In earlier stages of organogenesis, more regulon modules are enriched in the heart, spinal cord, and brain, and, transferred to cartilage and skeletal muscle in the later stages. During this process, regulations of brain development become more intensive and complex, as more regulons are enriched in the brain in PCW8.

Skeletal myogenesis starts early during development. It initiates within the somites following specifications of the premyogenic progenitors and skeletal myoblasts. A Human Skeletal Muscle Atlas indicated that the Muscle Score is relatively low in PCW3 and increased sharply from PCW4/5 ^73^. Our landscape indicated that the transcripts and regulons associated with skeletal muscle development and differentiation exhibit restricted patterns in somite (PCW3/4), and enrichment in myotome (PCW5), and followed by spreading patterns in skeletal muscle at PCW7 or later. The fetal liver, formed around PCW3/4 in human, becomes the major hematopoietic organ with the immigration of hematopoietic stem and progenitor cells at PCW6, then undergoes the hepatoblast-to-hepatocyte transition from PCW8 ^74, 75^. Hepatocytes are heterogeneous with their increasingly complex function during development ^76^. Our data suggested a key transition in the liver between PCW3-5 and PCW6- 8, where the enrichment of transcripts and regulons associated with cell proliferation and migration (PCW3-5) evolves into diversified functional genes such as hematopoiesis, blood coagulation, and lipid metabolism (PCW6-8). These spatiotemporal GRNs during human organogenesis would serve as a rich source for future studies on the fates of human cells. Meanwhile, understanding the developmental basis of organ formation and function helps pave the way for the elucidation and possible treatment of certain pathologies.

Congenital heart disease is the leading cause of human birth defects. During development, the human heart undergoes a series of complex morphogenetic processes that increase its ability to pump blood ^77^. The formation of trabeculae in the embryonic heart and the remodeling that occurs before birth is a conspicuous and challenging feature of human cardiogenesis. Abnormal trabeculae are a component of several cardiac pathologies and congenital heart disease ^78^. Here, trabecular architecture in the developing human embryo is analyzed and ETS1 is newly identified as a trabeculation-associated TF that modulates cardiomyocyte behavior during human cardiogenesis. Control of the rate and rhythm of atria and ventricle cardiomyocyte contractions is also essential for ensuring adequate blood circulation in the human body ^79^. The rhythmic contractions are triggered by the bioelectrical impulses intrinsically generated within the SAN. The molecular and cellular features of the SAN in the human developing heart that underpin its critical function are uncharted territory. Deficiency of Vsnl1, a core SAN cell cluster marker in mouse, not only reduces the beating rate of human induced pluripotent stem cell-derived cardiomyocytes but also the heart rate of mouse ^43^. Our SAN gene expression landscape shows human *VSNL1* is also expressed in the SAN region and knockdown of *vsnl1b* caused a reduction of the 2dpf embryos’ heart rate. We also found that *NR2F2*, a predicted Shox2 downstream gene in mouse ^80^, is co-expressed in the SAN region with SHOX2 and similar defects are observed in shox2 and nr2f2 deficient zebrafish embryos, which may explain the cardiac arrhythmias phenotype in patients with NR2F2 mutations ^81^. Future functional studies leveraging these substantial data will play a crucial role in improving our understanding of SAN and trabeculae development and function as well as translating these findings into tangible tools for the improved detection, prevention, and treatment of cardiac arrhythmias and abnormal trabeculae.

Neurogenesis occurs at an early stage with different temporal waves and is gradually followed by gliogenesis proceeding with spatial asynchrony. The construction of the human central nervous system requires precise coordination of numerous molecules and cells. Disorder in these processes will affect the structure and function of the central nervous system, thus causing neurological or psychiatric diseases. In past years, many studies provided single-cell atlas and spatial atlas of human and model organisms’ nervous system development ^21, 22, 26, 46, 52^. However, most of these studies focused on specific brain regions in the fetal period, and the research on the developmental whole brain in the human embryonic period remains limited. Our data contain transcriptional histograms with a spatial and temporal resolution of the complete nervous system of human embryos after gastrulation. During the embryonic stages, changes in the proportions of progenitors and neurons were evident (Fig. 4E and Fig. S9A-D). It is reported that excitatory neurons occur in pcd50-51 (PCW7-8) ^51, 82^ and the first GABA immunoreactive cell can be observed at the PCW6 ^82, 83^. Our data refines the time window for the origins of excitatory and inhibitory neurons and also indicated delayed neurogenesis of excitatory neurons in comparison with the inhibitory neurons. HMGA proteins confer the neurogenic potential of neocortical precursor cells by maintaining the open state of chromatin ^84^. We found that HMGA2 was widely expressed in PCW3-5 and specifically expressed in Pall VZ in PCW6-8. Regulon activity is also enriched only during PCW6-8. This indicates that HMGA2 plays an important role in the development of Pall VZ. Meanwhile, the physiological functions of the spinal cord are carried out by neural circuits comprised of molecularly distinct neuronal subtypes. The OLIG family plays a vital role in the development of the spinal cord. OLIG1 and OLIG2 promote the development of motor neurons in the ventral side of the spinal cord, and OLIG3 promotes the development of sensory neurons in the dorsal side of the spinal cord ^52^. Our OLIG family gene expression map shows that OLIG1 and OLIG2 are mainly in pMN, and OLIG3 is mainly in dp1. Meanwhile, we also found the presence of radial glial cells on the dorsal and ventral sides of the spinal cord (dp1, FP, and p3), and oligodendrocyte progenitor cells (pMN) and neural progenitor cells (dp6, p0) in the center. These all indicated that the spatial patterns of neuron subtypes helped the organized function of the spinal cord.

Although the placenta has a strong microbial defense mechanism to restrict vertical transmission during pregnancy, microorganisms that cause congenital diseases have likely evolved diverse mechanisms to bypass such defense. Both DNA and RNA viruses can traverse the maternal-fetal interface, causing congenital infection and disease ^85^. With the Stereo-seq data, we directly mapped the location of the receptors for 14 viruses in early embryonic tissues for the first time. The viral receptors are chronologically and spatially identified in different embryonic organs across PCW3-8, which could demonstrate the potential probability of ligand-receptor communications during early development in utero. Taken the hepatitis virus for example, as the World Health Organization pushes to eradicate HBV by 2030, the quest to halt perinatal mother-to-child transmission (MTCT) is becoming increasingly urgent ^86^. Antiviral therapy for mothers with a high viral load during pregnancy is of some significance to interrupt MTCT and ensure infant safety. However, the frequency which viruses pass from mothers to their offspring in utero remains inconclusive. In this study, although the identified cellular receptor NTCP for HBV is not enriched in the liver, the HBV and HCV receptor ASGR1 is expressed specifically in the liver across PCW3-8. Furthermore, the HCV receptors ASGR2 and CLDN1 expressed in the liver mostly after PCW5, indicating the importance of early pregnancy protection from hepatitis.

Vertical infection of SARS-CoV-2 during pregnancy remains controversial. Some previous studies demonstrated the capacity of SARS-CoV-2 to infect and propagate in the human placenta ^87, 88^, while others reported that the placenta minimally expresses the canonical cell-entry mediators for SARS-CoV-2, preventing fetal infection to some extent ^89, 90^. Single-cell RNA sequencing data from different fetal tissues showed that co-expression of ACE2 and TMPRSS2 in the intestine increased during gestation across the first and second trimester (PCW10-18), indicating that the fetal gastrointestinal tract is likely susceptible to SARS-CoV-2 infection due to exposures to potentially infected amniotic fluid^91^. It should be noted that all the early versions of SARS-CoV-2 relied on the cell receptor ACE2 binding to cells and the cellular enzyme TMPRSS2 breaking down part of the spike protein, thus revealing segments that allow the virus to fuse with human cells and immediately overload its RNA inside to make new viruses ^92^. In this study, the spatial co-expression of ACE2 and TMPRSS2 is found in the intestine epithelia from the time point of PCW8, but rarely co-expressed in the same cell, which may explain the previously referenced low infection rate from mother to fetus during early pregnancy. However, the Omicron variant carries a unique distinction from previous variants where its spike inefficiently utilizes the cellular protease TMPRSS2 that promotes cell entry via plasma membrane fusion. It instead places great dependency on cell entry on the endocytic pathway, indicating that the variant can enter cells for replication as long as it binds to ACE2 ^93, 94^. Therefore, the MTCT risk of women infected with Omicron or newer variants during pregnancy should be a concern in the future. Furthermore, women infected with Omicron or other variants during early pregnancy are susceptible to passing SARS-CoV-2 to the embryonic intestine tissue.

Genomic imprinting is typically involved in embryonic growth and development, and its dysregulation is associated with disorders in neurodevelopment and metabolic disease. Here we first show the spatiotemporal expression pattern of known imprinted genes and new monoallelic genes across multiple human embryos with spatial transcriptome data. The maternal-allelic expression pattern of PABPC1, a key component of the translational machinery, may explain the heterozygous mutation of this gene causing a developmental delay with impaired neurogenesis in cortical development ^95^. We also found several genes with tissue-unique patterns of parental-allelic expression in PCW4 and PCW7 embryos, which may reflect the genetic and epigenetic interaction on allelic gene expression during organogenesis. The identified function of some of these allelic-expressed genes, coupled with their chromosomal locus and tissue-specific pattern of expression during human organogenesis, hints at their possible involvement in various inherited disorders.

## Methods

### Ethics statement

Ethical approval of human embryo research was provided by the Ethics Committee of Obstetrics and Gynecology Hospital, Fudan University (2021- 121), and the Institutional Review Board of BGI-Shenzhen (BGI-IRB 22058). Human embryos were collected from women who voluntarily terminated the pregnancy at the Affiliated Obstetrics and Gynecology Hospital of Fudan University. The collection and use of human embryos were compliant with the current International Society for Stem Cell Research (ISSCR) Guidelines. All samples used in this study were not subjected to any other experiments. All procedures followed the ‘Interim Measures for the Administration of Human Genetic Resources’ administered by the Chinese Ministry of Health. After the termination of pregnancy, the patient was counseled by a senior clinician of this study explaining the nature of the research. Before the abortion procedure, each participant provided written informed consent for collection for research purposes.

### Embryo collection and preparation for Stereo-seq

2 to 3 embryos were collected at each stage during PCW3-8. Once obtained, human embryos were rinsed with PBS to remove surface blood. Embryos were then placed in a cryomold and embedded with optimal cutting temperature (OCT) compound (Sakura, 4583) within 30 minutes. Reagents in this experiment were prepared with sterilized water containing diethylpyrocarbonate (DEPC) (Sangon Biotech, B501005-0005), and all instruments were cleaned with sterilized water containing DEPC and RNAase Zap (Invitrogen, AM9780). Cryosections were performed in sagittal cut with a thickness of 10 μm using a Leika CM1950 cryostat. In each embryo, 1-10 sections with interval distance of 20-1800 µm (average interval distance: 310 µm) and covered the maximum sagittal section were obtained for Stereo-seq. An extra sagittal section next to each Stereo-seq section was also obtained for hematoxylin and eosin (H&E) staining or *in situ* hybridization (ISH) assays. Each section of Stereo-seq was then gently transferred to a Stereo-seq chip precooled at −20°C.

### Spatial transcriptome capture and sequencing for Stereo-seq

The spatial transcriptome of human embryo sections was obtained using the Stereo-seq technology, which relies on the DNBs that are photolithographically etched on the chip with 500-750 nm distance to capture tissue RNA ^22^. Spatial transcriptomics capture followed the previously described protocol ^22, 26^. The tissue section on the Stereo-seq chip was incubated at 37°C for 8 minutes, and fixed in methanol (Sigma, 34860) at −20°C for 40 minutes, followed by the staining of nuclei single-strand DNA (ssDNA) using nucleic acid dye (Thermo Fisher, Q10212) and imaging (Ti-7 Nikon Eclipse microscope). Sections were then permeabilized with 0.1% pepsin (Sigma, P7000) in 0.01M HCl buffer at 37°C for 12 minutes. The released RNAs were captured by DNBs on the Stereo-seq chip and reverse transcribed using SuperScript II (Invitrogen, 18064-014) at 42°C overnight. The chip was then washed with 0.1x SSC buffer (Thermo, AM9770) and digested with Tissue Removal buffer (10mM Tris-HCl, 25mM EDTA, 100mM NaCl, 0.5% SDS) at 37°C for 30 minutes. cDNAs were treated with Exonuclease I (NEB, M0293L) at 37°C for 1 hour, amplified with KAPA HiFi Hotstart Ready Mix (Roche, KK2602), purified using 0.6 × VAHTSTM DNA Clean Beads, and quantified using Qubit dsDNA HS assay kit (Invitrogen, Q32854). The amplified DNAs were fragmented with in-house Tn5 transposase, amplified with KAPA HiFi Hotstart Ready Mix, and purified with AMPure XP Beads. The qualified libraries were then sequenced on a DNBSEQ-Tx sequencer (MGI, China).

### Stereo-seq raw data processing

Stereo-seq raw data processing was performed as previously described ^22^. The coordinate identity (CID) sequences (read 1, 1-25 bp) were mapped to the coordinates of the chip, allowing 1 base mismatch. Molecular identity (MID) sequences (read 1, 26-35 bp) containing N bases or with a quality score below 10 for more than 2 bases were filtered out. cDNA sequences (read 2, 100 bp) were aligned to the reference genome (GRCh38) by STAR ^96^. The expression matrices with coordinates were generated from the above procedures.

### Cell segmentation

The nuclei ssDNA images were used to calculate the cell boundaries by the cell segmentation algorithm *spateo* ^97^ (https://github.com/aristoteleo/spateo-release). The grey images of nuclei ssDNA were converted to binary images using the calculated Gaussian-weighted threshold. The exact Euclidean distance transformation was performed to segment cell nuclei with overlapped regions. Labels representing different cell nuclei were transferred to pinpoint DNBs corresponding to spatial positions by the watershed algorithm (Roerdink and Meijster, 2000) with parameters of *distance=6, max_area=600*.

### Unsupervised spatially constrained clustering (SCC)

Unsupervised SCC clustering was conducted by *Scanpy* ^98^ with the bin50 count matrices and subsequently annotated the partitioned clusters at the organ level based on their highly differential genes. Count matrices were first normalized, and variable genes were identified by *Scanpy.* For each Stereo-seq chip, the 30-nearest neighbor graph based on gene expressions and the 8-nearest neighbor graph based on spatial coordinates were constructed by *Scanpy* ^98^ and *Squidpy* ^99^, respectively. A final unweighted neighborhood graph was computed by taking the union of expression and spatial connectivity, then used as input of the Leiden algorithm to identify clusters with tuned resolutions.

### Gene regulatory networks analysis

Gene regulatory networks (GRN) were inferred from Stereo-seq bin matrices according to the pySCENIC protocol ^100^. The databases used for pySCENIC were downloaded from https://resources.aertslab.org/cistarget/databases/. We used bin50 matrices (for embryos of PCW3-7) or bin200 matrices (for embryos of PCW8) to perform SCENIC analysis. Co-expression modules were inferred, pruned, and quantified from the count matrix by GRNBoost2, cisTarget, and AUCell, respectively. Together with the physical coordinates, regulon activity scores (RAS) ^101^ were visualized in the tissue context across developmental stages by *Scanpy*.

### Regulon downstream analysis

Spatially-dependent regulon modules were identified by Hotspot ^102^. First, the RAS matrix from pySCENIC was normalized by a negative binomial model (‘danb’). Then the k-nearest neighbor graph (10 nearest neighbors) was computed between bins based on spatial proximity. Autocorrelations were computed for each regulon. Lastly, spatially variable regulons were grouped as spatial modules with a minimum of 3 regulons per module and visualized by covariance heatmap. Bins with the top 5 percentile of module scores were considered as corresponding module locations. Sankey plots representing relationships between GRN modules and annotated organs were visualized by *Plotly*.

For each stage/chip, regulon specificity scores (RSS) were calculated across organs ^101^. RAS of each regulon were first normalized to a probability distribution. Organs were then represented as a vector of binary labels (target organ:1; others:0) and normalized to the probability distribution. These steps were followed by Jensen-Shannon Divergence (JSD) calculations to measure the differences between the two normalized probability distributions with a range from 0 to 1. Lastly, JSD scores were converted to the RSS scores using the formula: *RSS = 1 – sqrt(JSD)*. A higher RSS score indicates increased enrichment of regulon in the corresponding organ. We ranked regulons in each organ based on RSS scores and selected the top 5 or top 10 as organ-identity regulons. Heatmaps of the top 5 regulons across organs were visualized by *seaborn*.

### Data integration of multiple sections

The Python package *scanorama* was employed to integrate multiple sections and remove batch effects of multiple datasets ^103^. The UMI count matrices were normalized with *scanpy*. Integrated data were scaled and underwent *PCA* for dimension reduction. *Scanorama* was then used to correct batch effects, and dimensionality reduction was performed with *UMAP*. Leiden algorithm is used to identify clusters.

### Gene module analysis for cardiac development

The trabecular layer and compact layer were identified in the heart across PCW3-8 by calculating the gene module expression score using *scanpy.tl.score_genes* function with default parameters (ctrl_size=50, n_bins=25), with trabecular layer gene set (*CDKN1C*, *IRX3*, *BMP10*, *S1PR1*, *GJA5*, *ENG*, *FHL2*, *NOG*, *NOTCH1*, *TGFBR3*, *ETS1*) and compact layer gene set (*TBX20*, *HEY2*), respectively.

### Calculating cell-type percentage in brain regions and subregions

We selected 72 embryo sections with good morphology and data quality to extract the brain and spinal cord transcriptome data for re-clustering. Functional cell types including excitatory neurons, inhibitory neurons, radial glial cells, astrocytes, microglia, oligodendrocyte, oligodendrocyte progenitor cells, neural progenitor cells, and glia progenitor cells were annotated using previously reported markers (Table S6) ^20, 104–106^. Brain regions and subregions were annotated using previously reported markers ^47–50^. The proportion of a cell type (e.g. excitatory neurons) in a brain region (e.g. Fb) is calculated as the bin50 number of the cell type (e.g. excitatory neurons) divided by the total bin50 number of the brain region (e.g. Fb).

### Cell-type deconvolution for spatial transcriptomics

We initially performed cell segmentation for cell type mapping analysis in the spinal cord and gut as described above. Then, we downloaded the scRNA-seq data from the public database and aligned it to our spatial atlas using *Tangram*^107^. The scRNA-seq data of human embryonic spinal cord data was downloaded from GSE171892 ^52^ and the gut data of PCW7.9-8.4 was extracted from the fetal gut cell atlas ^58^ for annotation.

### Analysis of genes associated with virus infection and human developmental disorders

Virus receptors and genes for monogenic disease from the DDG2P database (V3.0) ^59^ were analyzed. After filtering out low-quality reads (genes count < 1), a list of 51 virus receptors and 1909 genes associated with developmental diseases remained, of which the expression levels in the selected organs at each developmental stage were aggregated, scaled, and further visualized by R package ComplexHeatmap (V2.12.0) ^108^. The DDG2P genes were clustered based on the scaled matrix, and the maximum method was adopted for the distance measure. After clustering, 1225 genes showed organ and developmental stage enrichment and were kept for final visualization.

### Cross-species comparisons between human and mouse

For the interspecies analysis, the spatial transcriptome data of mouse embryos from E9.5 to E14.5 were retrieved from the MOSTA database ^22^. At each developmental stage, the transcriptome data of three sections of human and mouse embryos were compared. The information on embryo sections used for comparison is shown in Table S14. The count matrices for all sections were processed in four steps: (1) obtain the orthologous gene in human and mouse from Ensembl ^109^ and mapped the homologous gene symbols from mouse to human by two R packages, biomaRt (V2.52.0) ^110^ and homologene (V1.4.68.19.3.27) (<u>GitHub - oganm/homologene: An r package that works as a</u> <u>wrapper to homologene</u>), (2) filter out mouse genes not orthologous to human, (3) filter out the matrices based on parameters including the number of counts and features detected, the fraction of mitochondrial genes, and the percentage of ribosomal genes, (4) for each developmental stage, normalize and integrate the gene expression matrices of three sections by Seurat ^111^. Then, the rds objects of the two species were further integrated at each corresponding developmental stage, and subjected to clustering using the standard pipeline of Seurat.

To identify differentially expressed genes between human and mouse embryos in representative anatomic regions (brain, spinal cord, eye, ear, liver, heart, lung, gut, kidney) at each developmental stage, we adopted the FindMarkers function in Seurat with default parameters, and the normalized RNA assay data from integrated objects were used for calculations. Go enrichment analysis was performed by R package clusterProfiler (V4.4.1) ^112^.

### Integrating spatial transcriptome and genome for allelic expression analysis

The embryonic samples used in the allelic analysis are listed in Table S20. The duplicate-marked transcriptome BAM file of each chip from Stereo-seq raw data was utilized to undergo the following processes: tissue division (samtools, version 1.15.1), sorting (samtools), splitting N cigar reads (SplitNCigarReads, GATK, version 4.1.8.1), BQSR with known site from NCBI dbsnp and 1000G project (BQSR, GATK), variant detection (HaplotypeCaller, GATK) with parameters “--ERC GVCF” and “--do-not-run-physical-phasing”. The whole genome sequencing of the embryo and decidua from PCW4 E3 and PCW7 E3 were performed by DNBseq T1&T5 and then aligned to human genome hg38 though SOAPnuke (Default: “-n 0.1 -q 0.5 -l 12 -Q 2 -G 2 -M 2”) and BWA. The genomic variant detection flow is the same as the transcriptome variant detection. The GVCF files of the embryonic genome, combined with the GVCF files (GenotypeGVCFs, GATK) of the transcriptome of all targeted tissues or maternal genome from the same embryo, transformed into the embryo genome and tissue-specific expression VCF (EaTSE VCF) and the embryo and mother VCF (EaM VCF). These VCF files passed filtering (--genotypegvcfs-stand-call-conf 30, -window 35, --cluster 3, FS > 60.0, QD < 2.0, DP < 5) and were annotated in terms of ensGene, cosmic70, exac03, and clinvar_20190305 (hg38) provided by annovar ^113^.

### Third-generation sequencing and phasing for allelic expression analysis

To improve the phasing result in the difficult-to-map genome regions, third-generation nanopore sequencing for PCW4 E3 embryo was performed with PromethION by BGI. Sequencing data is base-identified by guppy5 and aligned to hg38 with minimap2 (2.24-r1122). The genomic variant detection and phasing were then performed with Clair3 (v0.1-r12) ^114^. PCW7 E3, additionally sequenced by Pacbio underwent the same pipeline as PCW4 E3. The high-quality (GQ>10, DP>4) embryonic heterozygous variant and maternal homozygous variant from the EaM VCF file were selected to generate the preliminary parental phasing table. With assistance from the phasing blocks of the third-generation phasing VCF, the embryonic heterozygous variants, corresponding to maternal heterozygous variants, were added to the final phasing table.

### Parental monoallelic expression extraction and plot

The high-quality embryonic heterozygous variant and homozygous expression in certain tissue were extracted into tissue-specific monoallelic expression tables. Variants outside the gene region were filtered and tagged with parental allele information in reference to the previous final parental phasing table. The top 100 confident monoallelic expression genes (the number of variant >5 or the number of single variant >500) in each selected tissue were chosen to conduct subsequent allelic expression analysis. Variants of the top 100 tissue-specific monoallelic expression genes excluding the intron region, were scanned by mpileup of samtools (version 1.15.1), and plotted in parental allelic expression pattern with spatial location by matplotlib package (3.5.2). Furthermore, the parental expression rate was counted by comparing the number of parents reads in the same tissue and plotted through ggplot2 (3.4.0).

### *In Situ* Hybridization (ISH) assays

ISH was performed on 10 μm cryosections with RNAScope 2.0 multiplex fluorescent reagent kit assay (Advanced Cell Diagnostics, 323100) following the manufacturer’s instructions. In situ probes against human *ALB* (600941-C1), *MYL7* (831731-C1), *PMEL* (410671-C1), *SHH* (600951-C1), *STMN2* (525211- C2), *LMX1A* (540661-C1), *MYH6* (555381-C1) and *MYH7* (508201-C2) were used in combination with RNAScope 2.0 multiplex fluorescent reagent kit for target detection. Tissue sections were fixed in 10% neutral formalin reagent (NBF) (Solarbio, G2161), dehydrated through an ethanol series, processed using standard pre-treatment conditions, and followed by incubation with target probe as per the RNAScope 2.0 multiplex fluorescent reagent kit assay protocol. TSA-plus fluorescein, Cy3, and Cy5 fluorophores were used at 1:1500 dilution for the manual assay. Slides were imaged by PANNORAMIC MIDI II (3D Histech, Hungary) and viewed by Slide Converter.

### *In vivo* functional validation of human heart conduction genes

Knockdown of *roraa*, *rorab*, *shox2*, *vsnl1b*, *nr2f2,* and *elapor2b* was performed in zebrafish embryos by CRISPR/Cas9-induced mutagenesis. 4-6 CRISPR target sites for each gene were designed with CRISPR (https://zlab.bio/guide-design-resources). The oligo pairs for each target are shown in Table S21. The single-guide RNAs (sgRNAs) were generated with the T7 RiboMAX Express Large Scale RNA Production System (Promega, P1320) and purified with the RNeasy Mini Kit (QIAGEN, 74104). Mixed sgRNAs for every gene (final concentration 200 ng/μL) and the Cas9 protein (500 ng/μL, Novoprotein, E365) were co-injected into one-cell-stage embryos (2 nL/embryo) using the MPPI-3 Pressure Injector (Smith-Root). The injected embryos were treated with PTU (0.003%; MACKLIN, N816213) from 24 hours post-fertilization (hpf) and the heart rate was recorded by Fluorescence confocal microscope (OLYMPUS, BX51WI) 2 days post-fertilization (dpf). And the primer probes for the target genes detection were listed in Table S22.

### Immunofluorescent staining of pacemaker cells

2 dpf *Tg(myl7:H2B-EGFP)* transgenic zebrafish embryos were fixed in 1% formaldehyde for 1 hour at room temperature, then washed 3 times in 1 x PBS. The fixed embryos were blocked in 1 x PBS containing 10% donkey serum (absin, abs935), 2 mg/ml BSA (Solarbio, A8020), and 0.2% saponin (sigma, SAE0073) for 1 hour at room temperature and then incubated in 200 μl of Islet-1 (Isl1) primary antibody (GeneTex, GTX128201, 1:500 dilution) and GFP-Tag (7G9) mouse mAb (Abmart, M20004S, 1:1000 dilution) overnight at 4°C. The embryos were then stained with Anti-rabbit Alexa 594 (Abcam, ab150084, 1:500 dilution) and anti-mouse Alexa 488 (Abcam, ab150113, 1:500 dilution) secondary antibodies for 2 hours in the dark at room temperature. Images were taken by NIKON AXR HD25 laser scanning confocal microscope and raw data was processed with NIS Elements AR and ImageJ (v2.9.0).

## Acknowledgments

We thank all participating patients for their kind contributions. We thank Linying Wang, Binbin Jiang, Yanan Zhang, and Yixi Chen for the experimental assistance. Special thanks also go to Longqi Liu, Shiping Liu, Ying Gu, Yuxiang Li, Yinqi Bai, Lei Han, Lifang Wang, Sha Liao, Ao Chen, Kailong Ma, Shuxia Cao, Xiaoming Li, and Ruiling Zhang for their valuable scientific advice and technical supports. This work was also supported by China National GenBank (CNGB).

## Funding

National Key Research and Development Plan (2022YFC2703500, 2021YFC2700603, 2022YFC3400400, 2022YFC2703600, 2022YFC2703803, 2022YFC2703001)

National Natural Science Foundation of China (82088102, 82171613, 82192873, 82171688, 82271722, 82192864)

CAMS Innovation Fund for Medical Sciences (2019-I2M-5-064)

Collaborative Innovation Program of Shanghai Municipal Health Commission (2020CXJQ01)

Clinical Research Plan of SHDC (SHDC2020CR1008A)

Shanghai Clinical Research Center for Gynecological Diseases

Shanghai Urogenital System Diseases Research Center

Shanghai Frontiers Science Research Center of Reproduction and Development.

## Author contributions

HH conceived the idea

HH, XX, GD, YG, HY, JP, and LZ made the study design

JP, YL, YG, GD, HY, XX, and HH supervised the work

JP and GD designed the sample collection protocol

HC, ZL, YC, YT, XY, GZ, and YX collected the samples with the help of JP, GD, QC, YG, NM, HL, XL, TZ, and CX

YG, YL, and XJ designed the spatial transcriptome experiments

YL, ZL, QL, MZ, SS, YZ, PM, QQ, BJ, JN, and ML performed bioinformatics analysis, statistical analysis, and result visualization with the help of YG, XJ, GD, JP, HY, GZ, YX, HX, YC, JS, HH, XL, YW, and CX

JP, GD, and YM conducted ISH experiments; QW and HY performed zebrafish validation experiments

JP, YL, LZ, QL, HC, MZ, GZ, and SS wrote the first draft of the manuscript; JP, YG, GD, HY XX, and HH revised and finalized the manuscript

## Data and materials availability

All data that support the findings of this study will be available on request from the corresponding author. And the data will be publicly available when the paper is published. The data that support the findings of this study have been deposited into CNGB Sequence Archive (CNSA) ^115^ of China National GeneBank DataBase (CNGBdb) ^116^.

## Supplementary Materials

Figs. S1 to S14

Tables S1 to S22

Movies S1 to S14

## Reference

1 Homsy, J. et al. De novo mutations in congenital heart disease with neurodevelopmental and other congenital anomalies. Science 350, 1262–1266, doi:10.1126/science.aac9396 (2015).

2 Chen, B. et al. Maternal inheritance of glucose intolerance via oocyte TET3 insufficiency. Nature 605, 761–766, doi:10.1038/s41586-022-04756-4 (2022).

3 Barnat, M. et al. Huntington’s disease alters human neurodevelopment. Science 369, 787–793, doi:10.1126/science.aax3338 (2020).

4 Sadler, T. W. Susceptible periods during embryogenesis of the heart and endocrine glands. Environ Health Perspect 108 Suppl 3, 555–561, doi:10.1289/ehp.00108s3555 (2000).

5 Yan, L. et al. Single-cell RNA-Seq profiling of human preimplantation embryos and embryonic stem cells. Nat Struct Mol Biol 20, 1131–1139, doi:10.1038/nsmb.2660 (2013).

6 Xue, Z. et al. Genetic programs in human and mouse early embryos revealed by single-cell RNA sequencing. Nature 500, 593–597, doi:10.1038/nature12364 (2013).

7 Li, L. et al. Single-cell multi-omics sequencing of human early embryos. Nature cell biology 20, 847–858, doi:10.1038/s41556-018-0123-2 (2018).

8 Petropoulos, S. et al. Single-Cell RNA-Seq Reveals Lineage and X Chromosome Dynamics in Human Preimplantation Embryos. Cell 165, 1012–1026, doi:10.1016/j.cell.2016.03.023 (2016).

9 O’Rahilly, R. Early human development and the chief sources of information on staged human embryos. European journal of obstetrics, gynecology, and reproductive biology 9, 273–280, doi:10.1016/0028-2243(79)90068-6 (1979).

10 Azkue, J. J. External surface anatomy of the postfolding human embryo: Computer-aided, three-dimensional reconstruction of printable digital specimens. J Anat 239, 1438–1451, doi:10.1111/joa.13514 (2021).

11 Cao, J. et al. A human cell atlas of fetal gene expression. Science 370, doi:10.1126/science.aba7721 (2020).

12 Xu, Y. et al. A single-cell transcriptome atlas profiles early organogenesis in human embryos. Nature cell biology 25, 604–615, doi:10.1038/s41556-023-01108-w (2023).

13 Gao, S. et al. Tracing the temporal-spatial transcriptome landscapes of the human fetal digestive tract using single-cell RNA-sequencing. Nature cell biology 20, 721–734, doi:10.1038/s41556-018-0105-4 (2018).

14 Fawkner-Corbett, D. et al. Spatiotemporal analysis of human intestinal development at single-cell resolution. Cell 184, 810–826.e823, doi:10.1016/j.cell.2020.12.016 (2021).

15 Asp, M. et al. A Spatiotemporal Organ-Wide Gene Expression and Cell Atlas of the Developing Human Heart. Cell 179, 1647–1660.e1619, doi:10.1016/j.cell.2019.11.025 (2019).

16 Hou, X. et al. Integrating Spatial Transcriptomics and Single-Cell RNA-seq Reveals the Gene Expression Profling of the Human Embryonic Liver. Front Cell Dev Biol 9, 652408, doi:10.3389/fcell.2021.652408 (2021).

17 Popescu, D. M. et al. Decoding human fetal liver haematopoiesis. Nature 574, 365–371, doi:10.1038/s41586-019-1652-y (2019).

18 He, P. et al. A human fetal lung cell atlas uncovers proximal-distal gradients of differentiation and key regulators of epithelial fates. Cell 185, 4841–4860.e4825, doi:10.1016/j.cell.2022.11.005 (2022).

19 Garcia-Alonso, L. et al. Single-cell roadmap of human gonadal development. Nature 607, 540–547, doi:10.1038/s41586-022-04918-4 (2022).

20 Fan, X. et al. Single-cell transcriptome analysis reveals cell lineage specification in temporal-spatial patterns in human cortical development. Sci Adv 6, eaaz2978, doi:10.1126/sciadv.aaz2978 (2020).

21 Aldinger, K. A. et al. Spatial and cell type transcriptional landscape of human cerebellar development. Nat Neurosci 24, 1163–1175, doi:10.1038/s41593-021-00872-y (2021).

22 Chen, A. et al. Spatiotemporal transcriptomic atlas of mouse organogenesis using DNA nanoball-patterned arrays. Cell 185, 1777–1792.e1721, doi:10.1016/j.cell.2022.04.003 (2022).

23 Xia, K. et al. The single-cell stereo-seq reveals region-specific cell subtypes and transcriptome profiling in Arabidopsis leaves. Dev Cell 57, 1299–1310.e1294, doi:10.1016/j.devcel.2022.04.011 (2022).

24 Liu, C. et al. Spatiotemporal mapping of gene expression landscapes and developmental trajectories during zebrafish embryogenesis. Dev Cell 57, 1284–1298.e1285, doi:10.1016/j.devcel.2022.04.009 (2022).

25 Wang, M. et al. High-resolution 3D spatiotemporal transcriptomic maps of developing Drosophila embryos and larvae. Dev Cell 57, 1271–1283.e1274, doi:10.1016/j.devcel.2022.04.006 (2022).

26 Wei, X. et al. Single-cell Stereo-seq reveals induced progenitor cells involved in axolotl brain regeneration. Science 377, eabp9444, doi:10.1126/science.abp9444 (2022).

27 Fan, Z. et al. An E2F5-TFDP1-BRG1 Complex Mediates Transcriptional Activation of MYCN in Hepatocytes. Frontiers in Cell and Developmental Biology 9, doi:10.3389/fcell.2021.742319 (2021).

28 Yasui, K., Okamoto, H., Arii, S. & Inazawa, J. Association of over-expressed TFDP1 with progression of hepatocellular carcinomas. J Hum Genet 48, 609–613, doi:10.1007/s10038-003-0086-3 (2003).

29 Zhao, C. & Dahlman-Wright, K. Liver X receptor in cholesterol metabolism. J Endocrinol 204, 233–240, doi:10.1677/JOE-09-0271 (2010).

30 Siatecka, M. & Bieker, J. J. The multifunctional role of EKLF/KLF1 during erythropoiesis. Blood 118, 2044–2054, doi:10.1182/blood-2011-03-331371 (2011).

31 Musa, J., Aynaud, M. M., Mirabeau, O., Delattre, O. & Grunewald, T. G. MYBL2 (B-Myb): a central regulator of cell proliferation, cell survival and differentiation involved in tumorigenesis. Cell Death Dis 8, e2895, doi:10.1038/cddis.2017.244 (2017).

32 Jankovic, V. et al. Id1 restrains myeloid commitment, maintaining the self-renewal capacity of hematopoietic stem cells. Proc Natl Acad Sci U S A 104, 1260–1265, doi:10.1073/pnas.0607894104 (2007).

33 Ji, M. et al. Id2 intrinsically regulates lymphoid and erythroid development via interaction with different target proteins. Blood 112, 1068–1077, doi:10.1182/blood-2008-01-133504 (2008).

34 Petryniak, M. A., Potter, G. B., Rowitch, D. H. & Rubenstein, J. L. Dlx1 and Dlx2 control neuronal versus oligodendroglial cell fate acquisition in the developing forebrain. Neuron 55, 417–433, doi:10.1016/j.neuron.2007.06.036 (2007).

35 Bernal, J. Thyroid hormone receptors in brain development and function. Nat Clin Pract Endocrinol Metab 3, 249–259, doi:10.1038/ncpendmet0424 (2007).

36 Desmaris, E. et al. DMRT5, DMRT3, and EMX2 Cooperatively Repress Gsx2 at the Pallium-Subpallium Boundary to Maintain Cortical Identity in Dorsal Telencephalic Progenitors. J Neurosci 38, 9105-9121, doi:10.1523/jneurosci.0375-18.2018 (2018).

37 Urquhart, J. E. et al. DMRTA2 (DMRT5) is mutated in a novel cortical brain malformation. Clin Genet 89, 724–727, doi:10.1111/cge.12734 (2016).

38 Cao, Y. et al. In Vivo Dissection of Chamber-Selective Enhancers Reveals Estrogen-Related Receptor as a Regulator of Ventricular Cardiomyocyte Identity. Circulation, doi:10.1161/circulationaha.122.061955 (2023).

39 Sakamoto, T. et al. A Critical Role for Estrogen-Related Receptor Signaling in Cardiac Maturation. Circ Res 126, 1685–1702, doi:10.1161/circresaha.119.316100 (2020).

40 Pei, L. et al. Dependence of hippocampal function on ERRgamma-regulated mitochondrial metabolism. Cell Metab 21, 628–636, doi:10.1016/j.cmet.2015.03.004 (2015).

41 Grego-Bessa, J. et al. Notch signaling is essential for ventricular chamber development. Dev Cell 12, 415–429, doi:10.1016/j.devcel.2006.12.011 (2007).

42 Espinoza-Lewis, R. A. et al. Shox2 is essential for the differentiation of cardiac pacemaker cells by repressing Nkx2-5. Dev Biol 327, 376–385, doi:10.1016/j.ydbio.2008.12.028 (2009).

43 Liang, D. et al. Cellular and molecular landscape of mammalian sinoatrial node revealed by single-cell RNA sequencing. Nat Commun 12, 287, doi:10.1038/s41467-020-20448-x (2021).

44 Lee, J. M., Kim, H. & Baek, S. H. Unraveling the physiological roles of retinoic acid receptor-related orphan receptor α. Exp Mol Med 53, 1278–1286, doi:10.1038/s12276-021-00679-8 (2021).

45 Peng, Y. et al. Inhibition of TGF-β/Smad3 Signaling Disrupts Cardiomyocyte Cell Cycle Progression and Epithelial-Mesenchymal Transition-Like Response During Ventricle Regeneration. Front Cell Dev Biol 9, 632372, doi:10.3389/fcell.2021.632372 (2021).

46 Werling, D. M. et al. Whole-Genome and RNA Sequencing Reveal Variation and Transcriptomic Coordination in the Developing Human Prefrontal Cortex. Cell Rep 31, 107489, doi:10.1016/j.celrep.2020.03.053 (2020).

47 Gui, Y. et al. Single-nuclei chromatin profiling of ventral midbrain reveals cell identity transcription factors and cell-type-specific gene regulatory variation. Epigenetics Chromatin 14, 43, doi:10.1186/s13072-021-00418-3 (2021).

48 Luo, Z. et al. Human fetal cerebellar cell atlas informs medulloblastoma origin and oncogenesis. Nature 612, 787–794, doi:10.1038/s41586-022-05487-2 (2022).

49 Gonçalves, C. S., Le Boiteux, E., Arnaud, P. & Costa, B. M. HOX gene cluster (de)regulation in brain: from neurodevelopment to malignant glial tumours. Cell Mol Life Sci 77, 3797–3821, doi:10.1007/s00018-020-03508-9 (2020).

50 Braun, E. et al. Comprehensive cell atlas of the first-trimester developing human brain. bioRxiv, 2022.2010.2024.513487, doi:10.1101/2022.10.24.513487 (2022).

51 Silbereis, J. C., Pochareddy, S., Zhu, Y., Li, M. & Sestan, N. The Cellular and Molecular Landscapes of the Developing Human Central Nervous System. Neuron 89, 248–268, doi:10.1016/j.neuron.2015.12.008 (2016).

52 Rayon, T., Maizels, R. J., Barrington, C. & Briscoe, J. Single-cell transcriptome profiling of the human developing spinal cord reveals a conserved genetic programme with human-specific features. Development 148, doi:10.1242/dev.199711 (2021).

53 Shukla, N. D., Tiwari, V. & Valyi-Nagy, T. Nectin-1-specific entry of herpes simplex virus 1 is sufficient for infection of the cornea and viral spread to the trigeminal ganglia. Molecular vision 18, 2711 (2012).

54 Yan, H. et al. Sodium taurocholate cotransporting polypeptide is a functional receptor for human hepatitis B and D virus. eLife 1, e00049, doi:10.7554/eLife.00049 (2012).

55 Vyas, A. K. et al. Placental expression of asialoglycoprotein receptor associated with Hepatitis B virus transmission from mother to child. Liver Int 38, 2149–2158, doi:10.1111/liv.13871 (2018).

56 Zhang, Q. et al. Exosomes derived from hepatitis B virus-infected hepatocytes promote liver fibrosis via miR-222/TFRC axis. Cell Biol Toxicol, doi:10.1007/s10565-021-09684-z (2022).

57 Zhou, L. et al. ACE2 and TMPRSS2 are expressed on the human ocular surface, suggesting susceptibility to SARS-CoV-2 infection. Ocul Surf 18, 537–544, doi:10.1016/j.jtos.2020.06.007 (2020).

58 Elmentaite, R. et al. Single-Cell Sequencing of Developing Human Gut Reveals Transcriptional Links to Childhood Crohn’s Disease. Dev Cell 55, 771–783.e775, doi:10.1016/j.devcel.2020.11.010 (2020).

59 Wright, C. F. et al. Genetic diagnosis of developmental disorders in the DDD study: a scalable analysis of genome-wide research data. Lancet 385, 1305–1314, doi:10.1016/s0140-6736(14)61705-0 (2015).

60 Schanze, I. et al. NFIB Haploinsufficiency Is Associated with Intellectual Disability and Macrocephaly. American journal of human genetics 103, 752–768, doi:10.1016/j.ajhg.2018.10.006 (2018).

61 Sin, Y. Y., Baron, G., Schulze, A. & Funk, C. D. Arginase-1 deficiency. Journal of Molecular Medicine 93, 1287–1296, doi:10.1007/s00109-015-1354-3 (2015).

62 Lee, B. H., Jin, H. Y., Kim, G. H., Choi, J. H. & Yoo, H. W. Argininemia presenting with progressive spastic diplegia. Pediatr Neurol 44, 218–220, doi:10.1016/j.pediatrneurol.2010.11.003 (2011).

63 Bos, F. L. et al. CCBE1 is essential for mammalian lymphatic vascular development and enhances the lymphangiogenic effect of vascular endothelial growth factor-C in vivo. Circ Res 109, 486–491, doi:10.1161/circresaha.111.250738 (2011).

64 Barlow, D. P. & Bartolomei, M. S. Genomic imprinting in mammals. Cold Spring Harb Perspect Biol 6, doi:10.1101/cshperspect.a018382 (2014).

65 Haniffa, M. et al. A roadmap for the Human Developmental Cell Atlas. Nature 597, 196–205, doi:10.1038/s41586-021-03620-1 (2021).

66 Matthews, K. R. & Moralí, D. National human embryo and embryoid research policies: a survey of 22 top research-intensive countries. Regen Med 15, 1905–1917, doi:10.2217/rme-2019-0138 (2020).

67 Han, X. et al. Construction of a human cell landscape at single-cell level. Nature 581, 303–309, doi:10.1038/s41586-020-2157-4 (2020).

68 Menon, R. et al. Single-cell analysis of progenitor cell dynamics and lineage specification in the human fetal kidney. Development 145, doi:10.1242/dev.164038 (2018).

69 Zhong, S. et al. A single-cell RNA-seq survey of the developmental landscape of the human prefrontal cortex. Nature 555, 524–528, doi:10.1038/nature25980 (2018).

70 Ramond, C. et al. Understanding human fetal pancreas development using subpopulation sorting, RNA sequencing and single-cell profiling. Development 145, doi:10.1242/dev.165480 (2018).

71 Xu, Y. et al. Single-cell transcriptome analysis reveals the dynamics of human immune cells during early fetal skin development. Cell Rep 36, 109524, doi:10.1016/j.celrep.2021.109524 (2021).

72 Wang, P. et al. Dissecting the Global Dynamic Molecular Profiles of Human Fetal Kidney Development by Single-Cell RNA Sequencing. Cell Rep 24, 3554–3567.e3553, doi:10.1016/j.celrep.2018.08.056 (2018).

73 Xi, H. et al. A Human Skeletal Muscle Atlas Identifies the Trajectories of Stem and Progenitor Cells across Development and from Human Pluripotent Stem Cells. Cell Stem Cell 27, 158–176.e110, doi:10.1016/j.stem.2020.04.017 (2020).

74 Gruppuso, P. A. & Sanders, J. A. Regulation of liver development: implications for liver biology across the lifespan. J Mol Endocrinol 56, R115–125, doi:10.1530/jme-15-0313 (2016).

75 Wang, X. et al. Comparative analysis of cell lineage differentiation during hepatogenesis in humans and mice at the single-cell transcriptome level. Cell Res 30, 1109–1126, doi:10.1038/s41422-020-0378-6 (2020).

76 Kyrmizi, I. et al. Plasticity and expanding complexity of the hepatic transcription factor network during liver development. Genes Dev 20, 2293–2305, doi:10.1101/gad.390906 (2006).

77 Buijtendijk, M. F. J., Barnett, P. & van den Hoff, M. J. B. Development of the human heart. Am J Med Genet C Semin Med Genet 184, 7–22, doi:10.1002/ajmg.c.31778 (2020).

78 Stöllberger, C., Finsterer, J. & Blazek, G. Left ventricular hypertrabeculation/noncompaction and association with additional cardiac abnormalities and neuromuscular disorders. Am J Cardiol 90, 899–902, doi:10.1016/s0002-9149(02)02723-6 (2002).

79 Park, D. S. & Fishman, G. I. The cardiac conduction system. Circulation 123, 904–915, doi:10.1161/circulationaha.110.942284 (2011).

80 Hoffmann, S. et al. Network-driven discovery yields new insight into Shox2-dependent cardiac rhythm control. Biochim Biophys Acta Gene Regul Mech 1864, 194702, doi:10.1016/j.bbagrm.2021.194702 (2021).

81 Bashamboo, A. et al. Loss Function of the Nuclear Receptor NR2F2, Encoding COUP-TF2, Causes Testis Development and Cardiac Defects in 46,XX Children. American journal of human genetics 102, 487-493, doi:10.1016/j.ajhg.2018.01.021 (2018).

82 Shi, Y. et al. Mouse and human share conserved transcriptional programs for interneuron development. Science 374, eabj6641, doi:10.1126/science.abj6641 (2021).

83 Zecevic, N. & Milosevic, A. Initial development of gamma-aminobutyric acid immunoreactivity in the human cerebral cortex. J Comp Neurol 380, 495–506, doi:10.1002/(sici)1096-9861(19970421)380:4<495::aid-cne6>3.0.co;2-x (1997).

84 Kishi, Y., Fujii, Y., Hirabayashi, Y. & Gotoh, Y. HMGA regulates the global chromatin state and neurogenic potential in neocortical precursor cells. Nat Neurosci 15, 1127–1133, doi:10.1038/nn.3165 (2012).

85 Megli, C. J. & Coyne, C. B. Infections at the maternal-fetal interface: an overview of pathogenesis and defence. Nat Rev Microbiol 20, 67–82, doi:10.1038/s41579-021-00610-y (2022).

86 Drew, L. How to stop mother-to-child transmission of hepatitis B. Nature 603, S50–s52, doi:10.1038/d41586-022-00814-z (2022).

87 Fahmi, A. et al. SARS-CoV-2 can infect and propagate in human placenta explants. Cell Rep Med 2, 100456, doi:10.1016/j.xcrm.2021.100456 (2021).

88 Faure-Bardon, V. et al. Protein expression of angiotensin-converting enzyme 2, a SARS-CoV-2-specific receptor, in fetal and placental tissues throughout gestation: new insight for perinatal counseling. Ultrasound Obstet Gynecol 57, 242–247, doi:10.1002/uog.22178 (2021).

89 Lü, M., Qiu, L., Jia, G., Guo, R. & Leng, Q. Single-cell expression profiles of ACE2 and TMPRSS2 reveals potential vertical transmission and fetus infection of SARS-CoV-2. Aging (Albany NY) 12, 19880–19897, doi:10.18632/aging.104015 (2020).

90 Pique-Regi, R. et al. Does the human placenta express the canonical cell entry mediators for SARS-CoV-2? Elife 9, doi:10.7554/eLife.58716 (2020).

91 Beesley, M. A. et al. COVID-19 and vertical transmission: assessing the expression of ACE2/TMPRSS2 in the human fetus and placenta to assess the risk of SARS-CoV-2 infection. Bjog 129, 256–266, doi:10.1111/1471-0528.16974 (2022).

92 Hoffmann, M. et al. SARS-CoV-2 Cell Entry Depends on ACE2 and TMPRSS2 and Is Blocked by a Clinically Proven Protease Inhibitor. Cell 181, 271–280.e278, doi:10.1016/j.cell.2020.02.052 (2020).

93 Shuai, H. et al. Attenuated replication and pathogenicity of SARS-CoV-2 B.1.1.529 Omicron. Nature 603, 693-699, doi:10.1038/s41586-022-04442-5 (2022).

94 Meng, B. et al. Altered TMPRSS2 usage by SARS-CoV-2 Omicron impacts infectivity and fusogenicity. Nature 603, 706–714, doi:10.1038/s41586-022-04474-x (2022).

95 Wegler, M. et al. De novo variants in the PABP domain of PABPC1 lead to developmental delay. Genet Med 24, 1761–1773, doi:10.1016/j.gim.2022.04.013 (2022).

96 Dobin, A. et al. STAR: ultrafast universal RNA-seq aligner. Bioinformatics 29, 15–21, doi:10.1093/bioinformatics/bts635 (2013).

97 Qiu, X. et al. Spateo: multidimensional spatiotemporal modeling of single-cell spatial transcriptomics. bioRxiv, 2022.2012.2007.519417, doi:10.1101/2022.12.07.519417 (2022).

98 Wolf, F. A., Angerer, P. & Theis, F. J. SCANPY: large-scale single-cell gene expression data analysis. Genome Biol 19, 15, doi:10.1186/s13059-017-1382-0 (2018).

99 Palla, G. et al. Squidpy: a scalable framework for spatial single cell analysis. bioRxiv, 2021.2002.2019.431994, doi:10.1101/2021.02.19.431994 (2021).

100 Aibar, S. et al. SCENIC: single-cell regulatory network inference and clustering. Nat Methods 14, 1083–1086, doi:10.1038/nmeth.4463 (2017).

101 Suo, S. et al. Revealing the Critical Regulators of Cell Identity in the Mouse Cell Atlas. Cell Rep 25, 1436–1445 e1433, doi:10.1016/j.celrep.2018.10.045 (2018).

102 DeTomaso, D. & Yosef, N. Hotspot identifies informative gene modules across modalities of single-cell genomics. Cell Syst 12, 446–456.e449, doi:10.1016/j.cels.2021.04.005 (2021).

103 Hie, B., Bryson, B. & Berger, B. Efficient integration of heterogeneous single-cell transcriptomes using Scanorama. Nat Biotechnol 37, 685–691, doi:10.1038/s41587-019-0113-3 (2019).

104 Fu, Y. et al. Heterogeneity of glial progenitor cells during the neurogenesis-to-gliogenesis switch in the developing human cerebral cortex. Cell Rep 34, 108788, doi:10.1016/j.celrep.2021.108788 (2021).

105 Hatakeyama, J. et al. Hes genes regulate size, shape and histogenesis of the nervous system by control of the timing of neural stem cell differentiation. Development 131, 5539–5550, doi:10.1242/dev.01436 (2004).

106 Fan, X. et al. Spatial transcriptomic survey of human embryonic cerebral cortex by single-cell RNA-seq analysis. Cell Res 28, 730–745, doi:10.1038/s41422-018-0053-3 (2018).

107 Biancalani, T. et al. Deep learning and alignment of spatially resolved single-cell transcriptomes with Tangram. Nat Methods 18, 1352–1362, doi:10.1038/s41592-021-01264-7 (2021).

108 Gu, Z. Complex heatmap visualization. iMeta 1, e43, doi:https://doi.org/10.1002/imt2.43 (2022).

109 Yates, A. D. et al. Ensembl 2020. *Nucleic Acids Res* **48**, D682–D688, doi:10.1093/nar/gkz966 (2020).

110 Durinck, S., Spellman, P. T., Birney, E. & Huber, W. Mapping identifiers for the integration of genomic datasets with the R/Bioconductor package biomaRt. Nat Protoc 4, 1184–1191, doi:10.1038/nprot.2009.97 (2009).

111 Butler, A., Hoffman, P., Smibert, P., Papalexi, E. & Satija, R. Integrating single-cell transcriptomic data across different conditions, technologies, and species. Nat Biotechnol 36, 411–420, doi:10.1038/nbt.4096 (2018).

112 Wu, T. et al. clusterProfiler 4.0: A universal enrichment tool for interpreting omics data. Innovation (Camb) 2, 100141, doi:10.1016/j.xinn.2021.100141 (2021).

113 Wang, K., Li, M. & Hakonarson, H. ANNOVAR: functional annotation of genetic variants from high-throughput sequencing data. Nucleic Acids Res 38, e164, doi:10.1093/nar/gkq603 (2010).

114 Zheng, Z. et al. Symphonizing pileup and full-alignment for deep learning-based long-read variant calling. Nature Computational Science 2, 797–803, doi:10.1038/s43588-022-00387-x (2022).

115 Guo, X. et al. CNSA: a data repository for archiving omics data. Database (Oxford) 2020, doi:10.1093/database/baaa055 (2020).

116 Chen, F. Z. et al. CNGBdb: China National GeneBank DataBase. Yi Chuan 42, 799–809, doi:10.16288/j.yczz.20-080 (2020).

