## Supplemental figures for "Spatiotemporal transcriptome atlas of human embryos after gastrulation"

**Affiliations**
[1] Obstetrics and Gynecology Hospital, Institute of Reproduction and Development, Fudan University, Shanghai, China
[2] BGI-Shenzhen, Shenzhen, China
[3] The Fourth Affiliated Hospital, International Institutes of Medicine, Zhejiang University School of Medicine, Yiwu, China
[4] Key Laboratory of Reproductive Genetics (Ministry of Education), Department of Reproductive Endocrinology, Women's Hospital, Zhejiang University School of Medicine, Hangzhou, China
[5] Research Units of Embryo Original Diseases, Chinese Academy of Medical Sciences (No. 2019RU056), Shanghai, China
[6] The International Peace Maternal and Child Health Hospital, School of Medicine, Shanghai Jiao Tong University, Shanghai, China
[7] The Third Affiliated Hospital of Zhengzhou University, Zhengzhou, China
[8] Hangzhou Women's Hospital, Hangzhou, China
[9] Medical Technology College of Hebei Medical University, Shijiazhuang, China.
[10] Shenzhen Engineering Laboratory for Birth Defects Screening, Shenzhen, China
[11] Guangdong Provincial Key Laboratory of Genome Read and Write, Shenzhen, China

### These authors contributed equally to this work.

**The PDF file includes:**

Figs. S1 to S14
Captions Tables for S1 to S22

**Other Supplementary Materials for this manuscript include the following:**

Movies S1 to S14

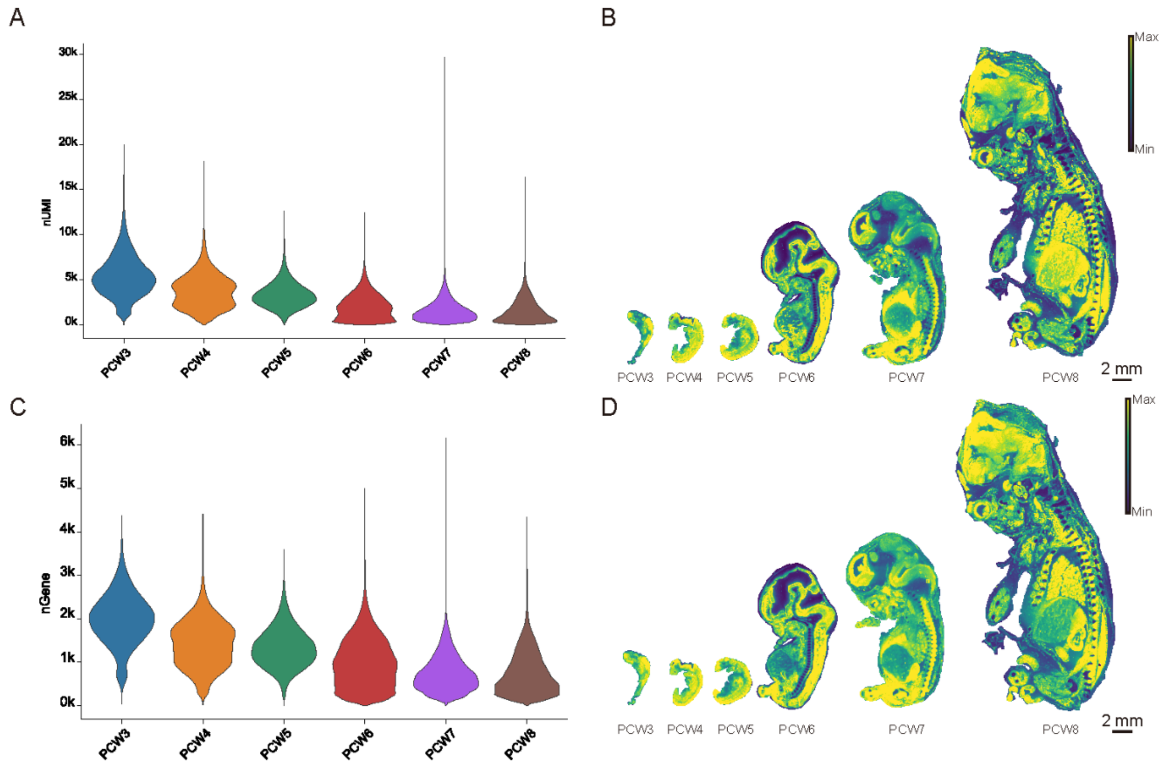


**Fig. S1. The count number of UMIs and genes across stages.** (A) Violin plot showing the count number of UMIs (bin50) in each stage. (B) Spatial visualization of the count number of UMIs (bin50) in each stage. (C) Violin plot showing the count number of genes (bin50) in each stage. (D) Spatial visualization of the count number of genes (bin50) in each stage.

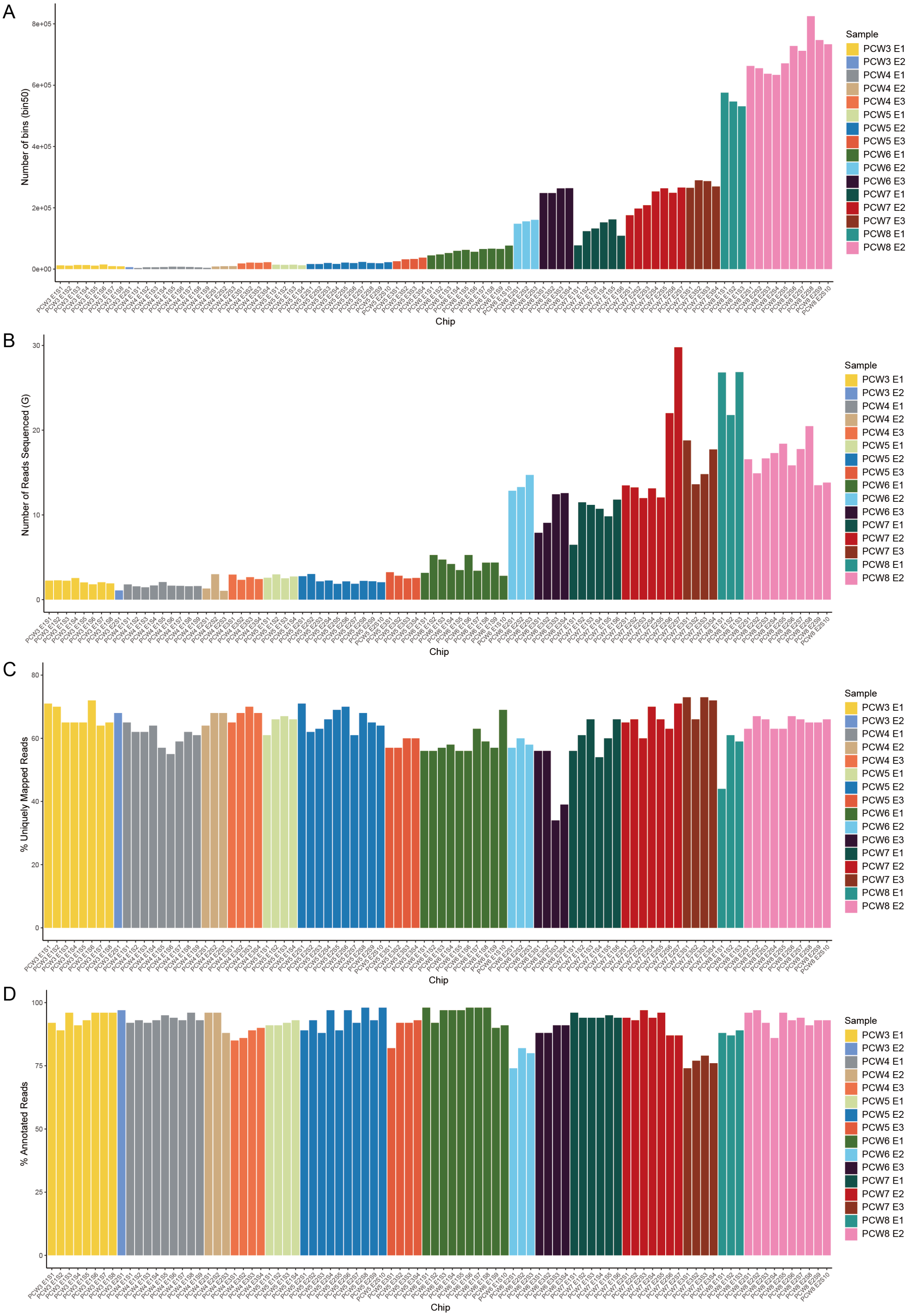
A
Number of bins (bin50)
8e+05
6e+05
4e+05
2e+05
0e+00
Chip
Sample
PCW3 E1
PCW3 E2
PCW4 E1
PCW4 E2
PCW4 E3
PCW5 E1
PCW5 E2
PCW5 E3
PCW6 E1
PCW6 E2
PCW6 E3
PCW7 E1
PCW7 E2
PCW7 E3
PCW8 E1
PCW8 E2
B
Number of Reads Sequenced (G)
30
20
10
0
Chip
C
% Uniquely Mapped Reads
80
60
40
20
0
Chip
D
% Annotated Reads
100
75
50
25
0
Chip

**Fig. S2. Quality control of Stereo-seq data.** (A) Barplot showing the number of bins (bin50) for all chips. (B) Barplot showing the number of reads sequenced for all chips. (C) Barplot showing the ratio of uniquely mapping reads for all chips. (D) Barplot showing the ratio of annotated reads for all chips.

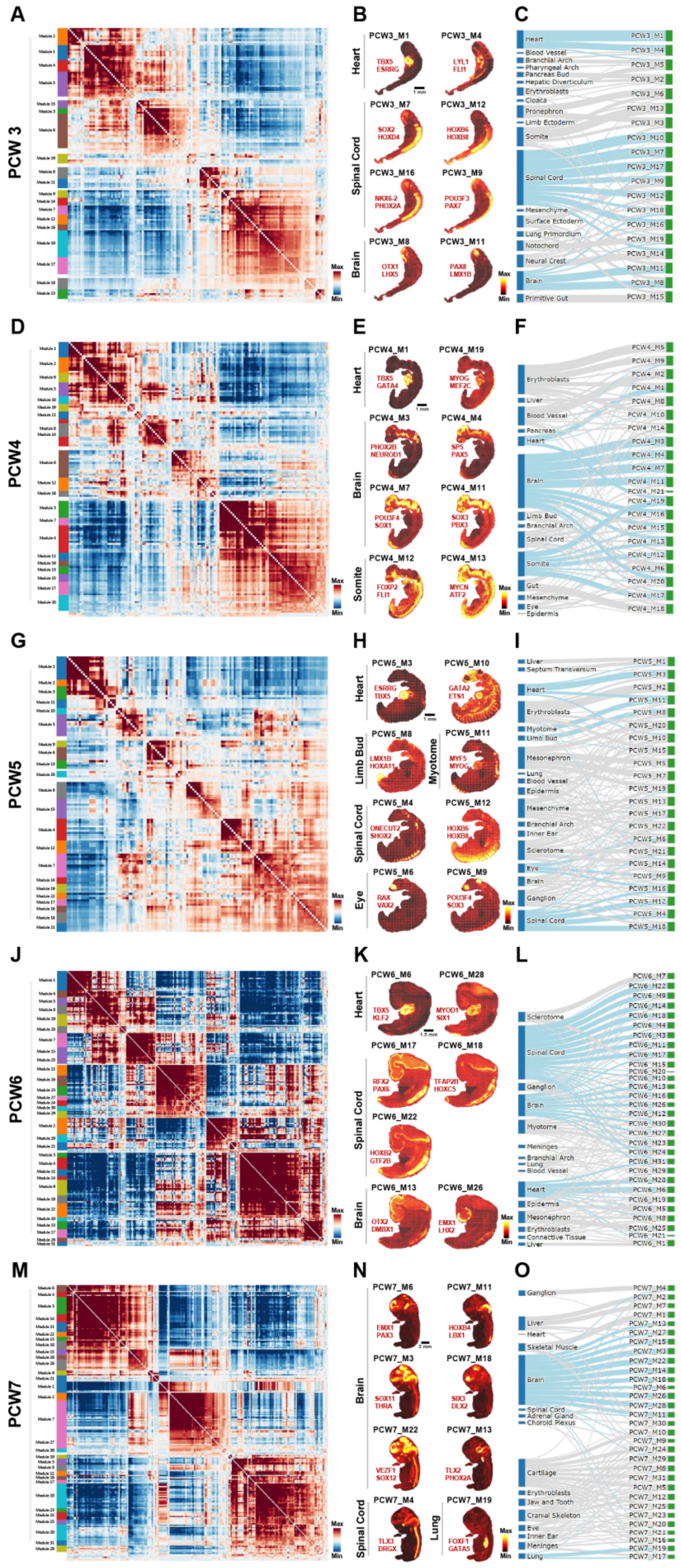

A
B
C
D
E
F
G
H
I
J
K
L
M
N
O
PCW 3
PCW4
PCW5
PCW6
PCW7
Heart
Spinal Cord
Brain
Somite
Limb Bud
Myotome
Eye
Lung
Max
Min

**Fig. S3. Regulon modules identified in main fetal organs based on the activity scores and spatial coordinates across PCW3-7.** (A, D, G, J, M) Regulon modules grouped by Hotspot based on pairwise local correlation at PCW3, PCW4, PCW5, PCW6, and PCW7, respectively. (B, E, H, K, N) Spatial patterns of regulon modules in representative organs at PCW3, PCW4, PCW5, PCW6, and PCW7, respectively. (C, F, I, L, O) Sankey plot showing the relationship between developmental organs and regulon modules at PCW3, PCW4, PCW5, PCW6, and PCW7, respectively. Representative organs are highlighted in blue.

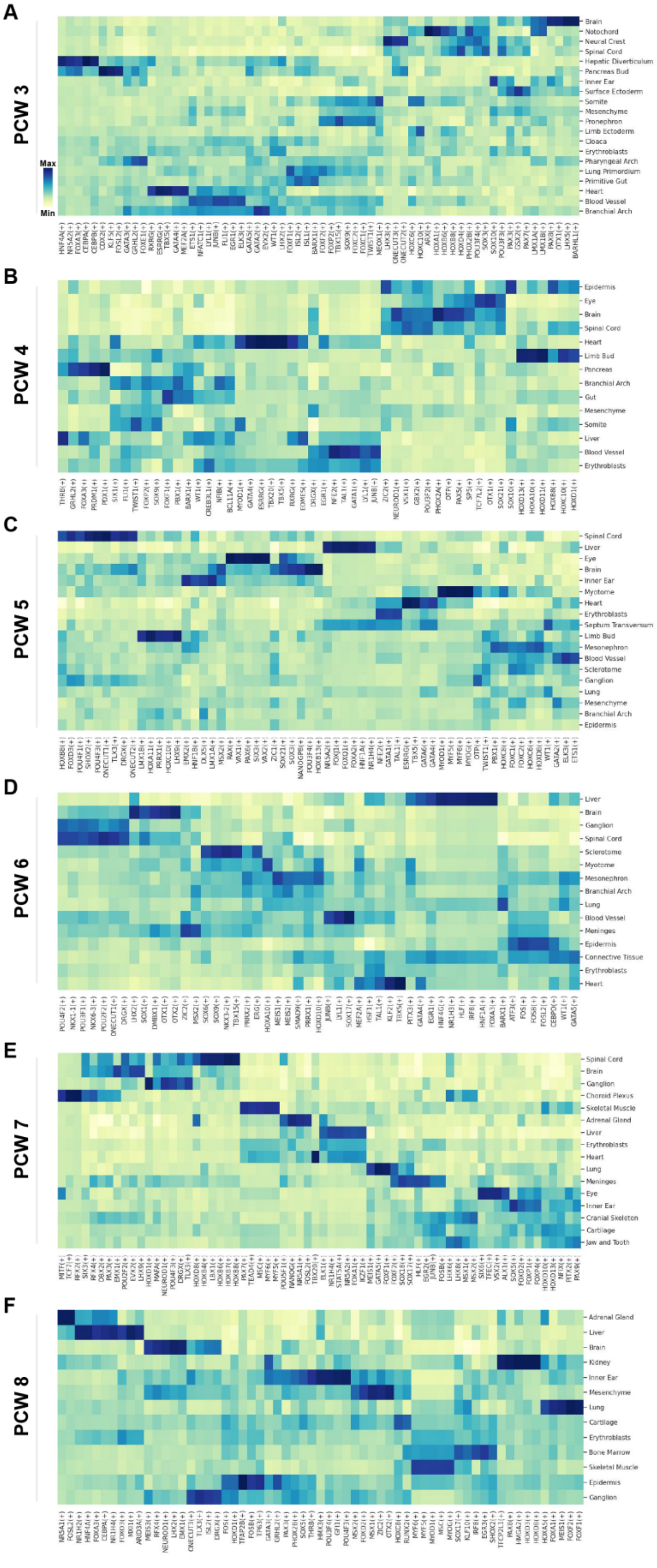
A
PCW 3
Max
Min
Brain
Notochord
Neural Crest
Spinal Cord
Hepatic Diverticulum
Pancreas Bud
Inner Ear
Surface Ectoderm
Somite
Mesenchyme
Pronephron
Limb Ectoderm
Cloaca
Erythroblasts
Pharyngeal Arch
Lung Primordium
Primitive Gut
Heart
Blood Vessel
Branchial Arch
B
PCW 4
Epidermis
Eye
Brain
Spinal Cord
Heart
Limb Bud
Pancreas
Branchial Arch
Gut
Mesenchyme
Somite
Liver
Blood Vessel
Erythroblasts
C
PCW 5
Spinal Cord
Liver
Eye
Brain
Inner Ear
Myotome
Heart
Erythroblasts
Septum Transversum
Limb Bud
Mesonephron
Blood Vessel
Sclerotome
Ganglion
Lung
Mesenchyme
Branchial Arch
Epidermis
D
PCW 6
Liver
Brain
Ganglion
Spinal Cord
Sclerotome
Myotome
Mesonephron
Branchial Arch
Lung
Blood Vessel
Meninges
Epidermis
Connective Tissue
Erythroblasts
Heart
E
PCW 7
Spinal Cord
Brain
Ganglion
Choroid Plexus
Skeletal Muscle
Adrenal Gland
Liver
Erythroblasts
Heart
Lung
Meninges
Eye
Inner Ear
Cranial Skeleton
Cartilage
Jaw and Tooth
F
PCW 8
Adrenal Gland
Liver
Brain
Kidney
Inner Ear
Mesenchyme
Lung
Cartilage
Erythroblasts
Bone Marrow
Skeletal Muscle
Epidermis
Ganglion

**Fig. S4. Organ-specific regulons across human developmental stages.** (A-F) Heatmaps showing the top5 organ-specific regulons based on RSS across PCW3 to PCW8.

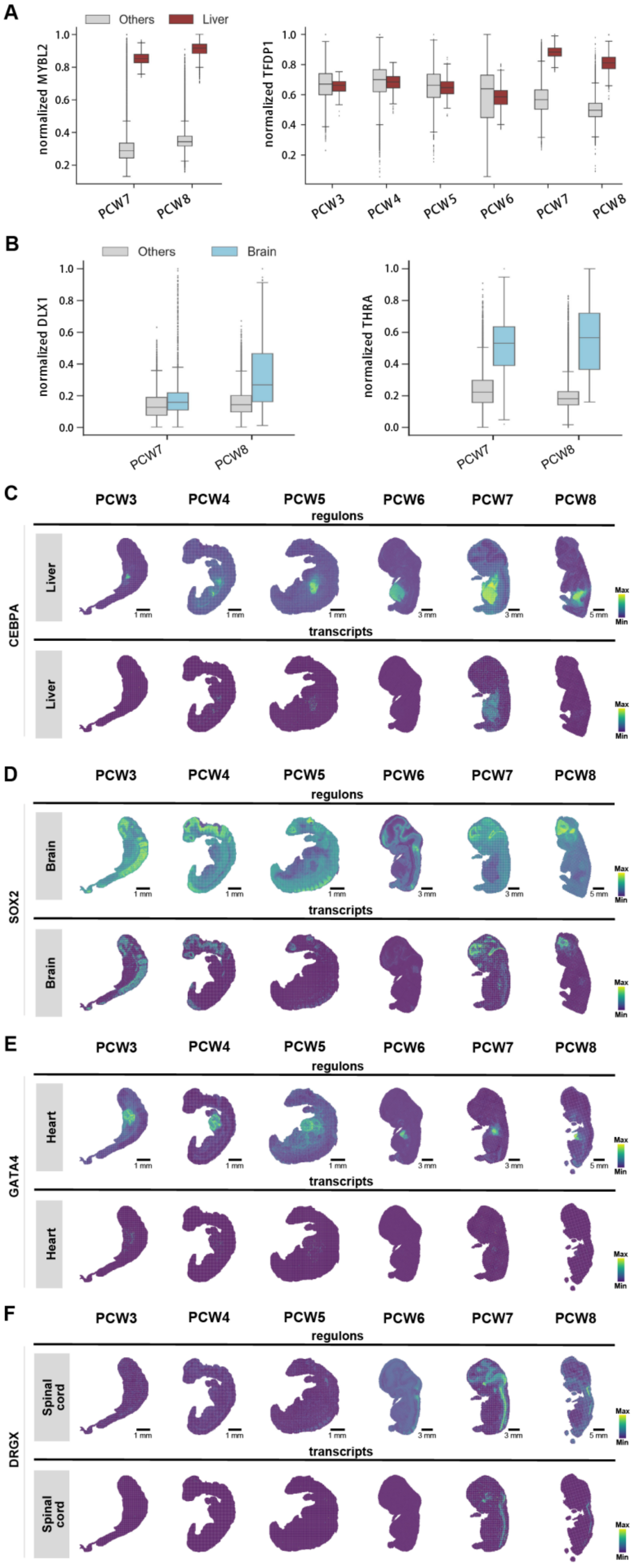
A
Others
Liver
normalized MYBL2
normalized TFDP1
PCW3
PCW4
PCW5
PCW6
PCW7
PCW8
B
Others
Brain
normalized DLX1
normalized THRA
C
regulons
transcripts
Liver
CEBPA
D
Brain
SOX2
E
Heart
GATA4
F
Spinal cord
DRGX
Max
Min

**Fig. S5. Spatial patterns of organ-specific regulons.** (A) Boxplot showing the normalized regulon activity of MYBL2 and TFDP1 within liver and other organs from PCW3 to PCW8. (B) Boxplot showing the normalized regulon activity of DLX1 and THRA within brain and other organs from PCW3 to PCW8. (C-F) Spatial visualization pattern of CEBPA, SOX2, GATA4, DRGX regulon activity and transcript expressions from PCW3 to PCW8.

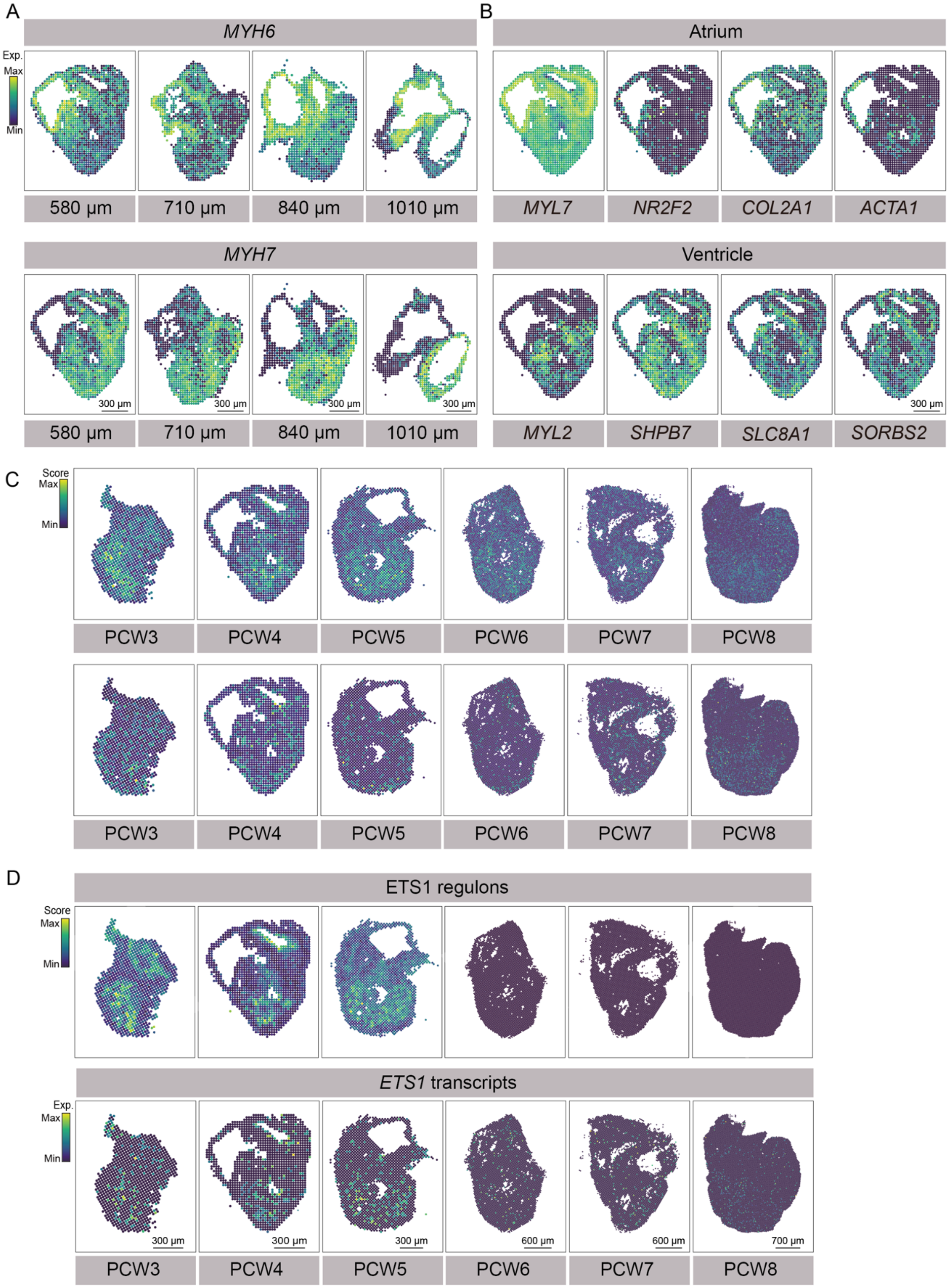


**Fig. S6. Marker gene expression and spatial visualization of trabecular modules in the heart.** (A) The atrium-specific gene *MYH6* and ventricle-specific gene *MYH7* were spatially visualized in the serial sections of PCW4 heart, showcasing their localization.

(B) Beyond *MYH6* and *MYH7*, additional atrium-specific genes *MYL7*, *COL2A1*, *NR2F2* and *ACTA1*, and ventricle-specific genes *MYL2*, *SLC8A1*, *HSPB7* and *SORBS2* were also visually mapped in the PCW4 heart. (C) Gene modules for the trabecular layer (upper) and compact layer (lower) were spatially visualized in the heart. (D) Spatial visualization of *ETS1* expression and regulon activity in heart across PCW3-8.

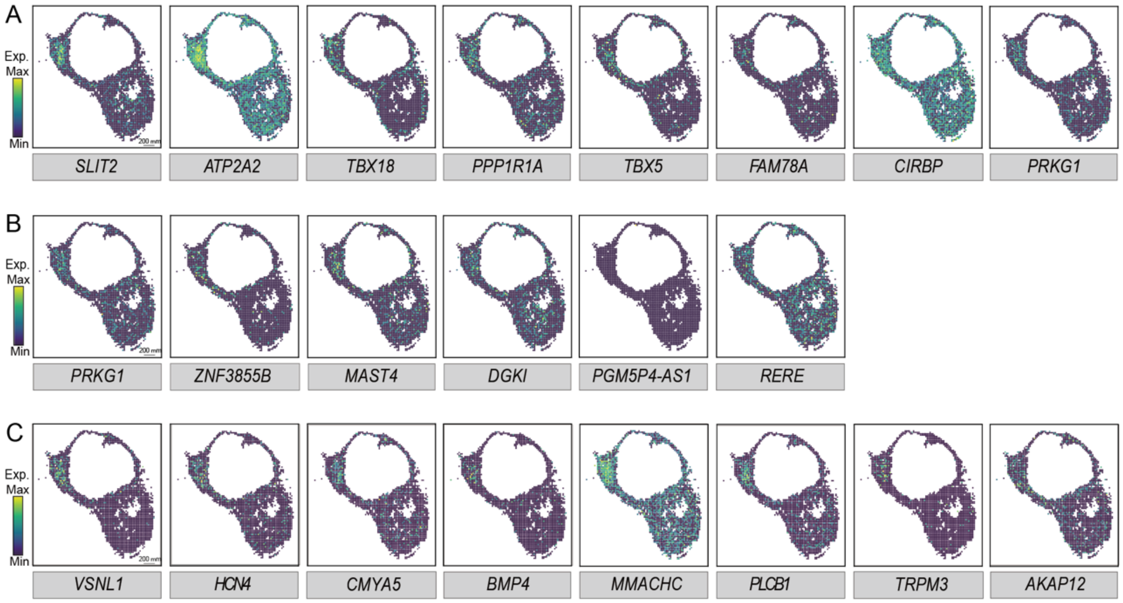


**Fig. S7. The spatial visualization of downstream genes of SHOX2, RORA and NR2F2 at the PCW6 heart.** (A) Spatial visualization of the gene expression of SHOX2 targets in PCW6 heart. (B) Spatial visualization of the gene expression of RORA targets in PCW6 heart. (C) Spatial visualization of *NR2F2*-colocalized target genes in PCW6 heart, including classic pacemaker cell markers *VSNL1* and *HCN4*.

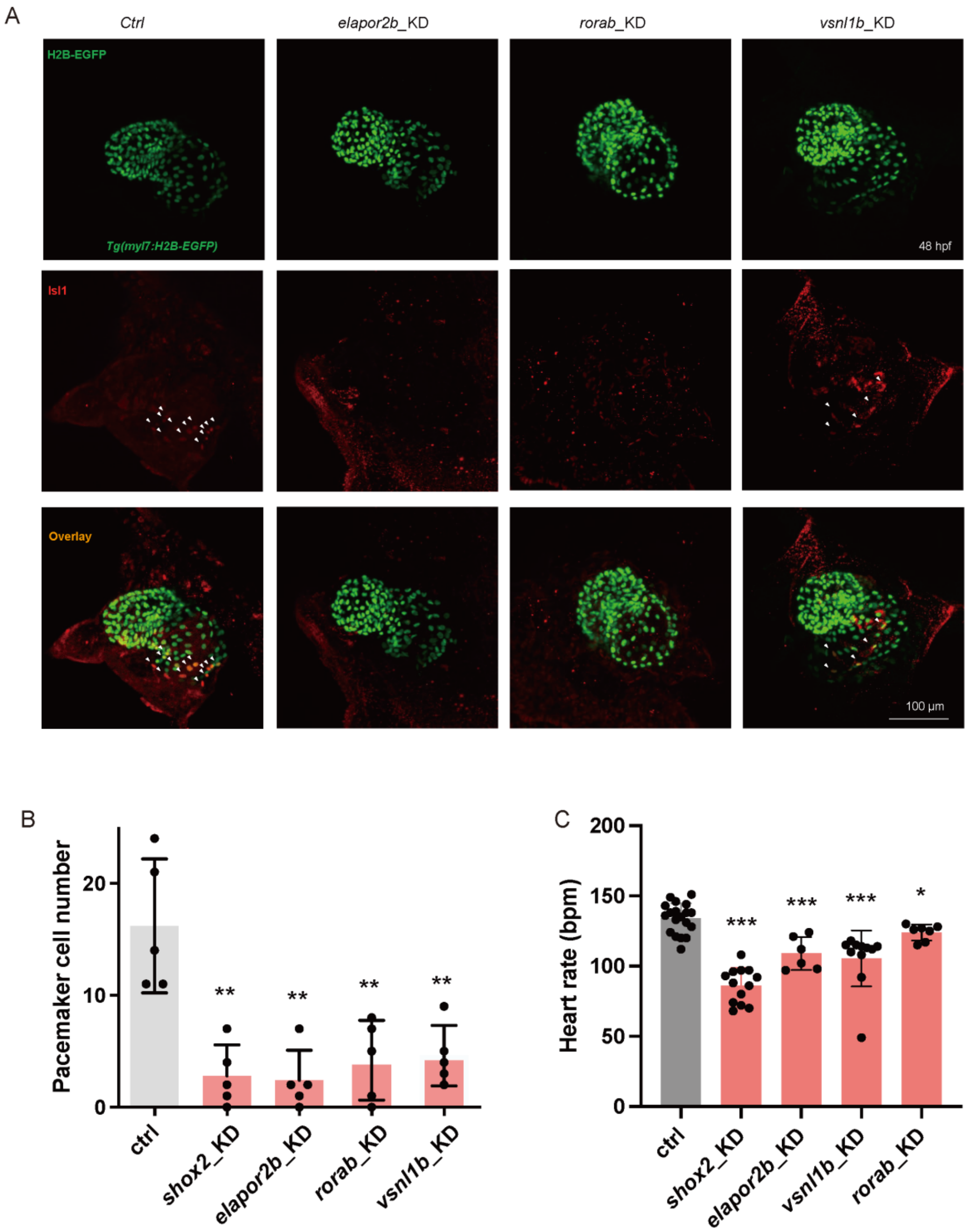


**Fig. S8.**
**Functional validation of *elapor2b* (*KIAA1324L* ortholog), *rorab* and *vsnl1b* in zebrafish.** (A) Immunohistochemistry of SAN in zebrafish at 48 hpf. Pacemaker cell nuclei are Isl1+ (red) in*Tg(myl7:H2B-EGFP)* transgenic fish (denoted by white arrows). (B) The pacemaker cell number in control, *shox2*, *elapor2b, rorab* and *vsnl1b* knockdown zebrafish at 48 hpf. **$p < 0.01$. (C) The heart rate of control, *shox2*, *elapor2b, rorab* and *vsnl1b* knockdown zebrafish at 48 hpf. *$p < 0.05$, ***$p < 0.001$.

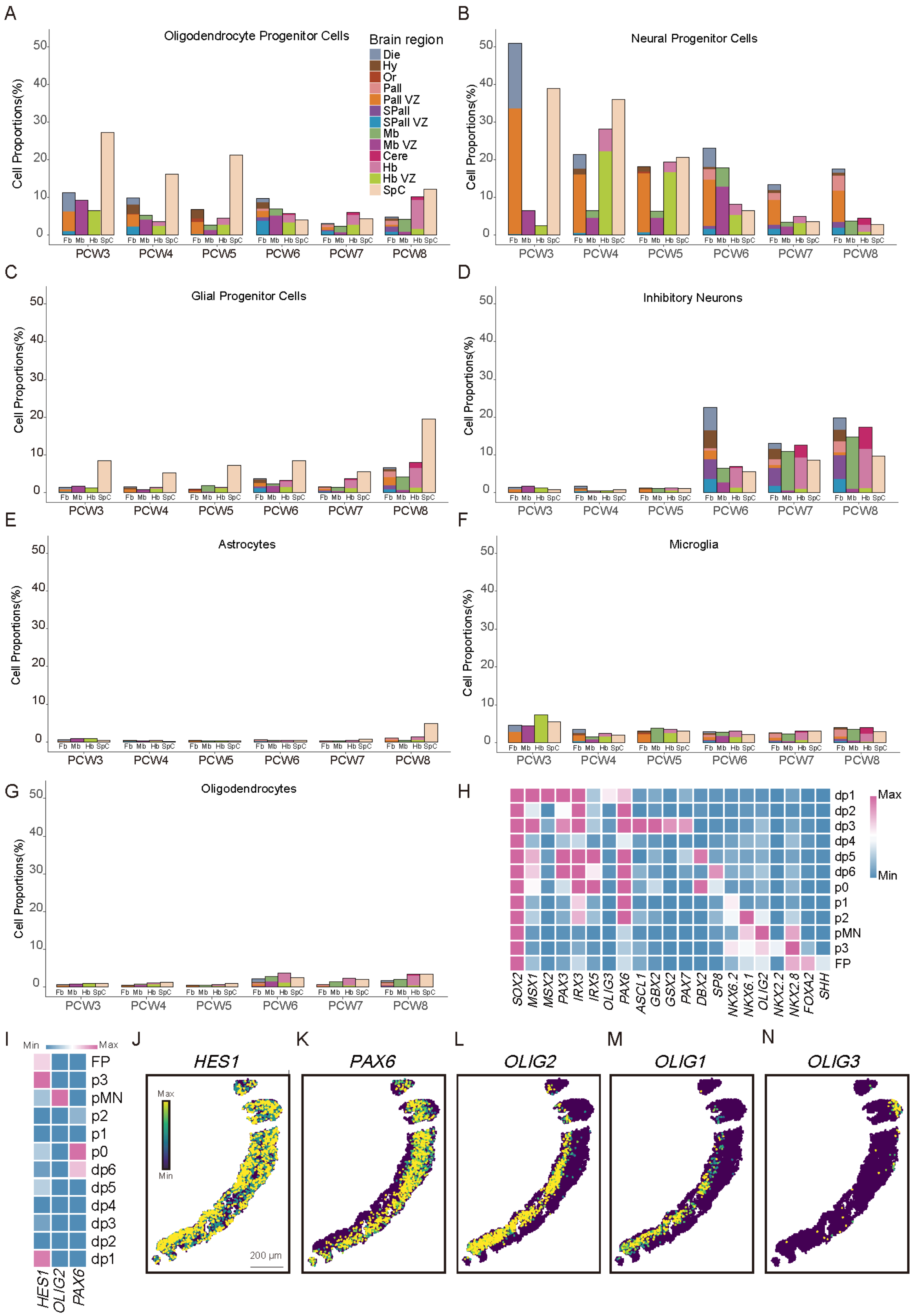

A
Oligodendrocyte Progenitor Cells
Brain region
Die
Hy
Or
Pall
Pall VZ
SPall
SPall VZ
Mb
Mb VZ
Cere
Hb
Hb VZ
SpC
B
Neural Progenitor Cells
C
Glial Progenitor Cells
D
Inhibitory Neurons
E
Astrocytes
F
Microglia
G
Oligodendrocytes
Cell Proportions(%)
Fb Mb Hb SpC
PCW3 PCW4 PCW5 PCW6 PCW7 PCW8
H
dp1 dp2 dp3 dp4 dp5 dp6 p0 p1 p2 pMN p3 FP
Max
Min
SOX2 MSX1 MSX2 PAX3 IRX3 IRX5 OLIG3 PAX6 ASCL1 GBX2 GSX2 PAX7 DBX2 SP8 NKX6.2 NKX6.1 OLIG2 NKX2.2 NKX2.8 FOXA2 SHH
I
Min Max
FP p3 pMN p2 p1 p0 dp6 dp5 dp4 dp3 dp2 dp1
HES1 OLIG2 PAX6
J
HES1
Max
Min
200 μm
K
PAX6
L
OLIG2
M
OLIG1
N
OLIG3

**Fig. S9. Cellular proportions of nerve-associated cells in the embryonic central nervous system.** (A-G) Barplot showing the cell proportion of oligodendrocyte progenitor cells, neural progenitor cells, glial progenitor cells, inhibitory neurons, astrocytes, microglia and oligodendrocytes in each brain region. (H) Heatmap showing the expression of markers used to identify dorsoventral (DV) domains in human spinal cord of PCW3. (I) Heatmap showing the expression of *HES1*, *PAX6* and *OLIG2* of anatomical structures in spinal cord at PCW3. (J-N) Spatial visualization of the expression patterns of *HES1*, *PAX6*, *OLIG2*, *OLIG1*, *OLIG3* in spinal cord based on single-cell segmentation.

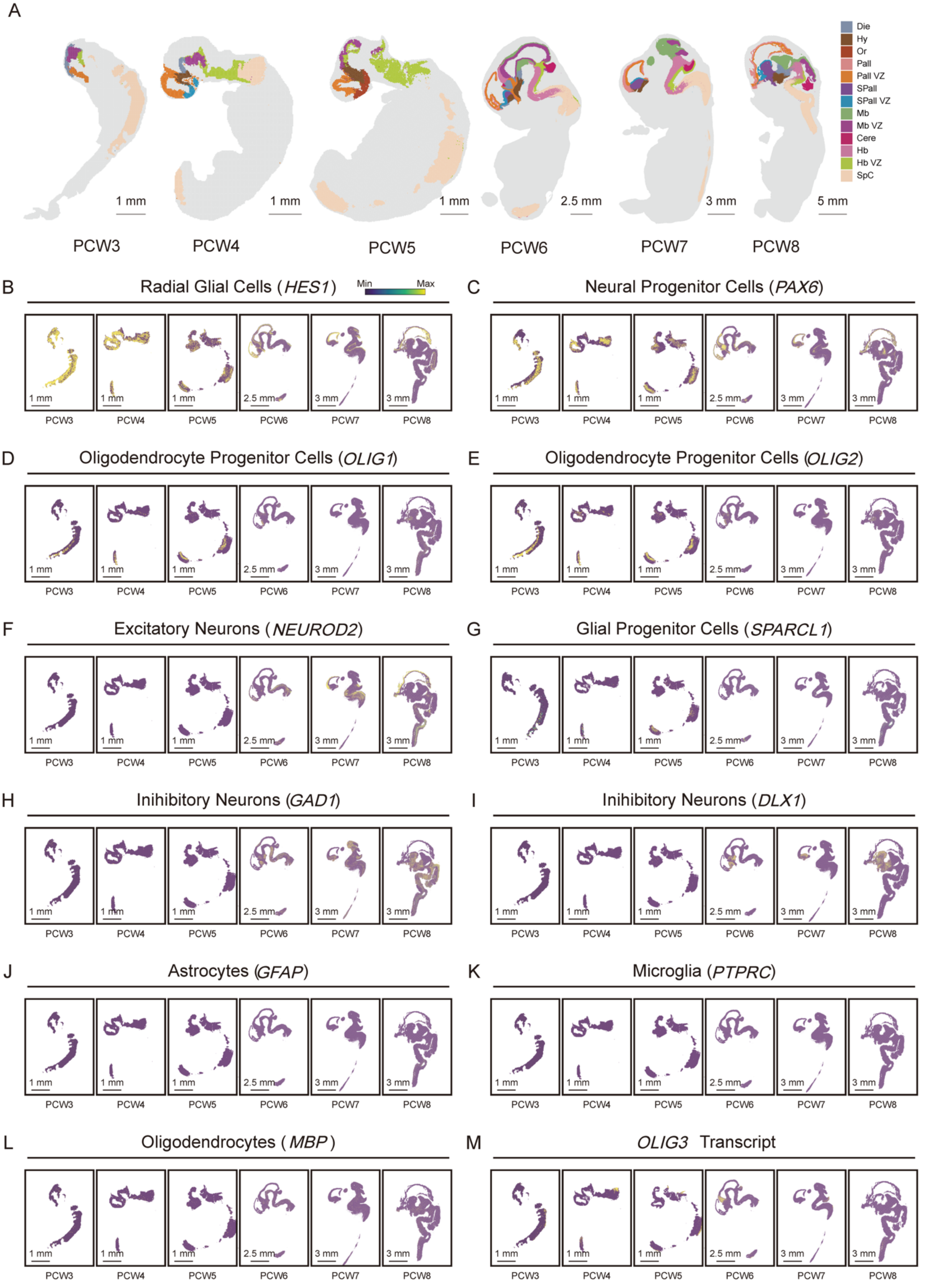
A
Die
Hy
Or
Pall
Pall VZ
SPall
SPall VZ
Mb
Mb VZ
Cere
Hb
Hb VZ
SpC
1 mm
1 mm
1 mm
2.5 mm
3 mm
5 mm
PCW3
PCW4
PCW5
PCW6
PCW7
PCW8
B
Radial Glial Cells (HES1)
Min
Max
C
Neural Progenitor Cells (PAX6)
D
Oligodendrocyte Progenitor Cells (OLIG1)
E
Oligodendrocyte Progenitor Cells (OLIG2)
F
Excitatory Neurons (NEUROD2)
G
Glial Progenitor Cells (SPARCL1)
H
Inihibitory Neurons (GAD1)
I
Inihibitory Neurons (DLX1)
J
Astrocytes (GFAP)
K
Microglia (PTPRC)
L
Oligodendrocytes (MBP)
M
OLIG3 Transcript

**Fig. S10. The spatial pattern of the marker gene expression.** (A) Spatiotemporal transcriptomic atlas of human embryonic nervous system. (B-M) Spatial visualization of gene expression for *HES1*, *PAX6*, *OLIG1*, *OLIG2*, *NEUROD2*, *SPARCL1*, *GAD1*, *DLX1*, *GFAP*, *PTPRC*, *MBP*, *OLIG3* in human embryonic nervous system.

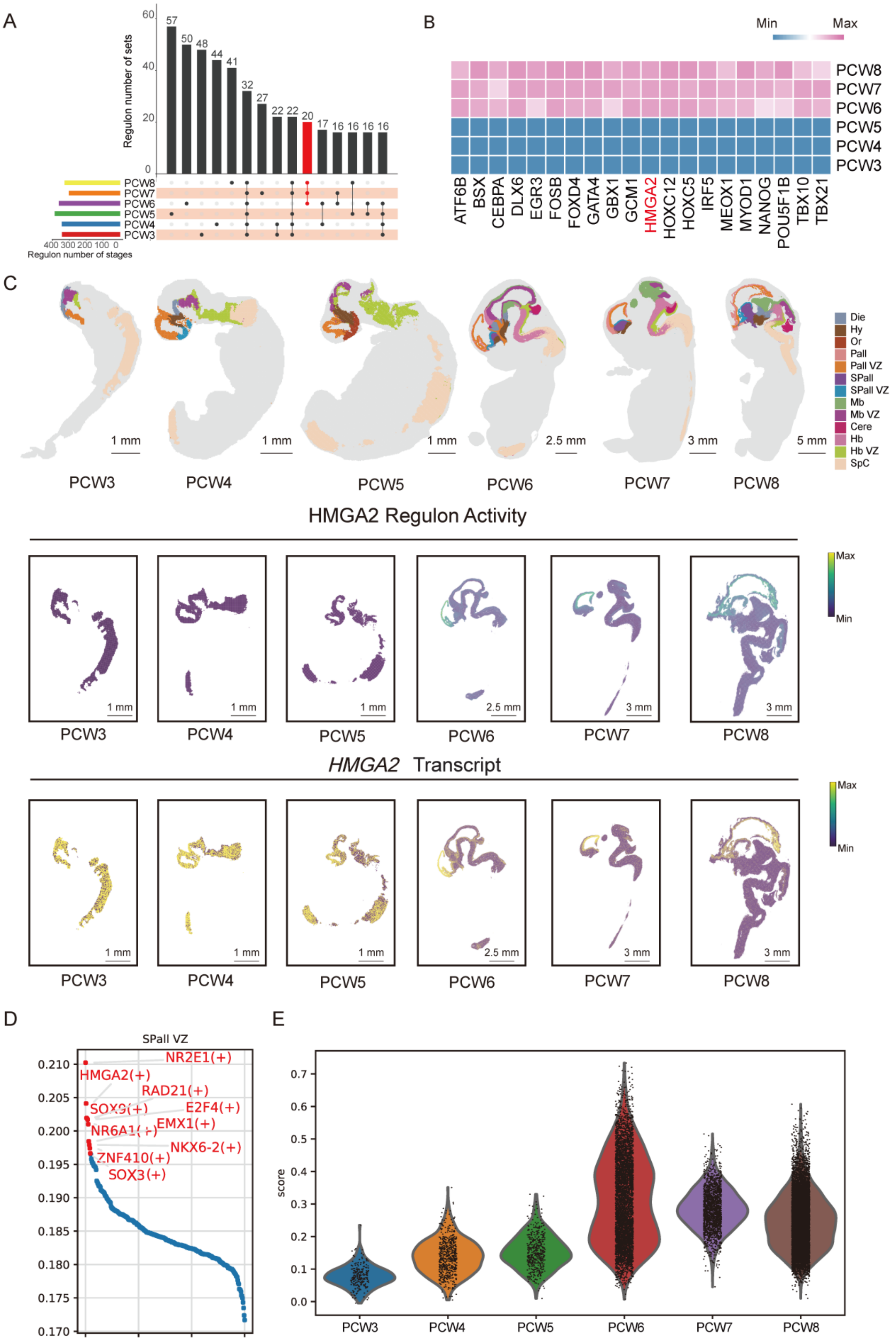
A
Regulon number of sets
Regulon number of stages
PCW8
PCW7
PCW6
PCW5
PCW4
PCW3
B
Min
Max
ATF6B
BSX
CEBPA
DLX6
EGR3
FOSB
FOXD4
GATA4
GBX1
GCM1
HMGA2
HOXC12
HOXC5
IRF5
MEOX1
MYOD1
NANOG
POU5F1B
TBX10
TBX21
C
Die
Hy
Or
Pall
Pall VZ
SPall
SPall VZ
Mb
Mb VZ
Cere
Hb
Hb VZ
SpC
HMGA2 Regulon Activity
HMGA2 Transcript
D
SPall VZ
NR2E1(+)
HMGA2(+)
RAD21(+)
E2F4(+)
EMX1(+)
NKX6-2(+)
ZNF410(+)
SOX3(+)
E
score

**Fig. S11. Temporal dynamics of regulon activity in central nervous system development.** (A) Upset plot showing regulon brain region at each stage. (B) Heatmap showing regulon activity at each stage. (C) Spatial visualization pattern of the HMGA2 regulon activity and expression. (D) Top 10 regulons of SPall VZ. (E) Violin plot showing gene set score of the HMGA2 target genes.

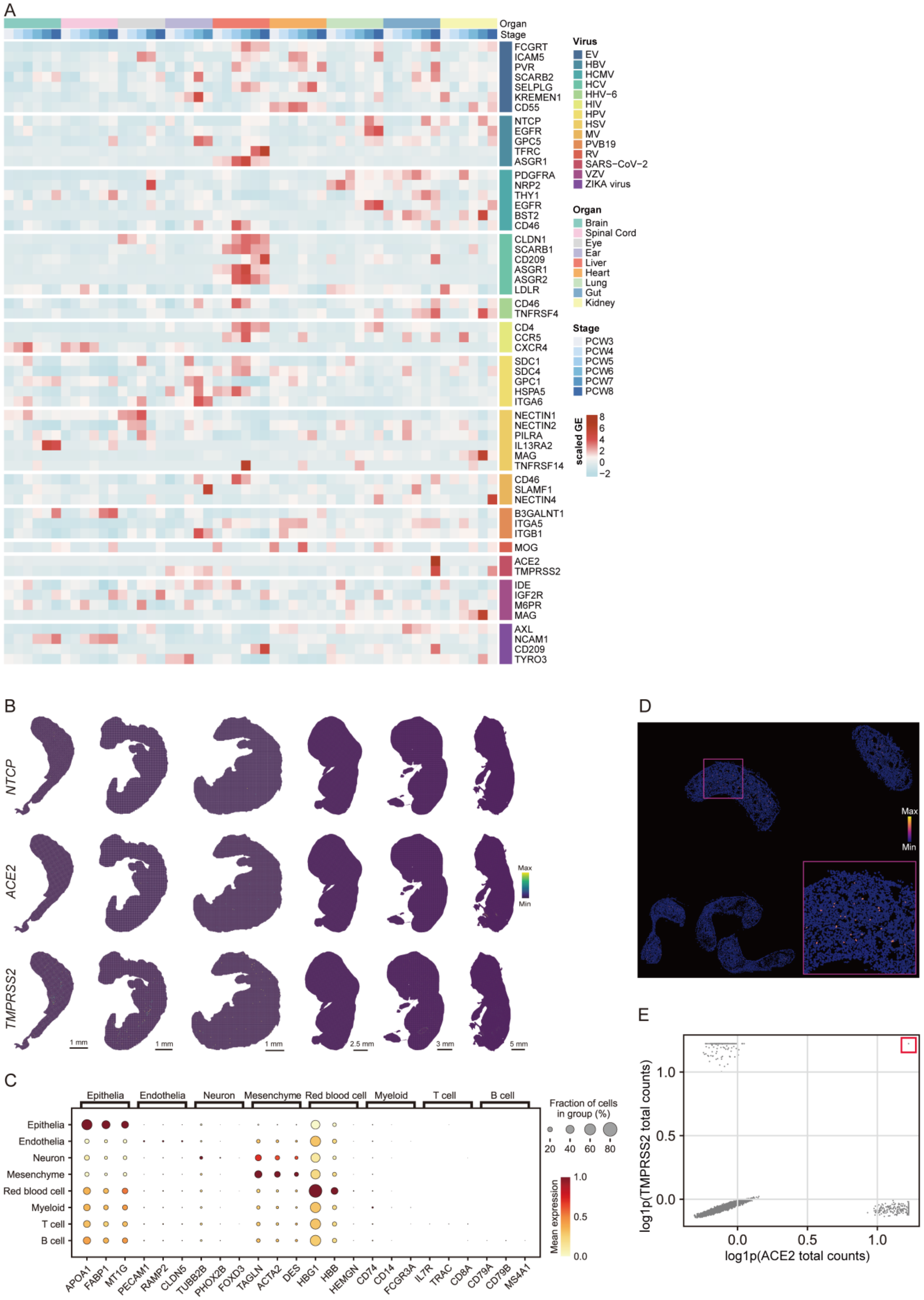

A
Organ
Stage
FCGRT
ICAM5
PVR
SCARB2
SELPLG
KREMEN1
CD55
NTCP
EGFR
GPC5
TFRC
ASGR1
PDGFRA
NRP2
THY1
EGFR
BST2
CD46
CLDN1
SCARB1
CD209
ASGR1
ASGR2
LDLR
CD46
TNFRSF4
CD4
CCR5
CXCR4
SDC1
SDC4
GPC1
HSPA5
ITGA6
NECTIN1
NECTIN2
PILRA
IL13RA2
MAG
TNFRSF14
CD46
SLAMF1
NECTIN4
B3GALNT1
ITGA5
ITGB1
MOG
ACE2
TMPRSS2
IDE
IGF2R
M6PR
MAG
AXL
NCAM1
CD209
TYRO3
Virus
EV
HBV
HCMV
HCV
HHV−6
HIV
HPV
HSV
MV
PVB19
RV
SARS−CoV−2
VZV
ZIKA virus
Organ
Brain
Spinal Cord
Eye
Ear
Liver
Heart
Lung
Gut
Kidney
Stage
PCW3
PCW4
PCW5
PCW6
PCW7
PCW8
scaled GE
8
6
4
2
0
−2
B
NTCP
ACE2
TMPRSS2
Max
Min
1 mm
1 mm
1 mm
2.5 mm
3 mm
5 mm
C
Epithelia
Endothelia
Neuron
Mesenchyme
Red blood cell
Myeloid
T cell
B cell
Epithelia
Endothelia
Neuron
Mesenchyme
Red blood cell
Myeloid
T cell
B cell
APOA1
FABP1
MT1G
PECAM1
RAMP2
CLDN5
TUBB2B
PHOX2B
FOXD3
TAGLN
ACTA2
DES
HBG1
HBB
HEMGN
CD74
CD14
FCGR3A
IL7R
TRAC
CD8A
CD79A
CD79B
MS4A1
Fraction of cells
in group (%)
20
40
60
80
Mean expression
1.0
0.5
0.0
D
Max
Min
E
log1p(TMPRSS2 total counts)
1.0
0.5
0.0
0.0
0.5
1.0
log1p(ACE2 total counts)

**Fig. S12. Organ and cell type vulnerability to viruses.** (A) Heatmap showing the normalized expression level of receptors for 14 viruses, including enterovirus (EV), hepatitis B virus (HBV), cytomegalovirus (HCMV), hepatitis C virus (HCV), human herpesvirus 6 (HHV-6), human immunodeficiency virus (HIV), human papillomavirus (HPV), herpes simplex virus (HSV), measles morbillivirus (MV), parvovirus B19 (PVB19), rubella virus (RV), severe acute respiratory syndrome coronavirus 2 (SARS-CoV-2), varicella zoster virus (VZV), and ZIKA virus in 9 representative anatomic regions (brain, spinal cord, eye, ear, liver, heart, lung, gut, kidney) in embryo sections from PCW3 to 8. Embryo sections are same as those in Fig. 5A. (B) Spatial visualization of the expression patterns of *NTCP*/*ACE2*/*TMPRSS2* expression from PCW3 to PCW8. (C) Bubbleplot showing the cell markers expression among all cell types. (D) Spatial visualization of *TMPRSS2* expression based on single-cell segmentation in gut at PCW8. (E) Scatterplot showing the co-expression pattern of *ACE2* and *TMPRSS2* expression within the same cell base on single-cell-segmented gut at PCW8.

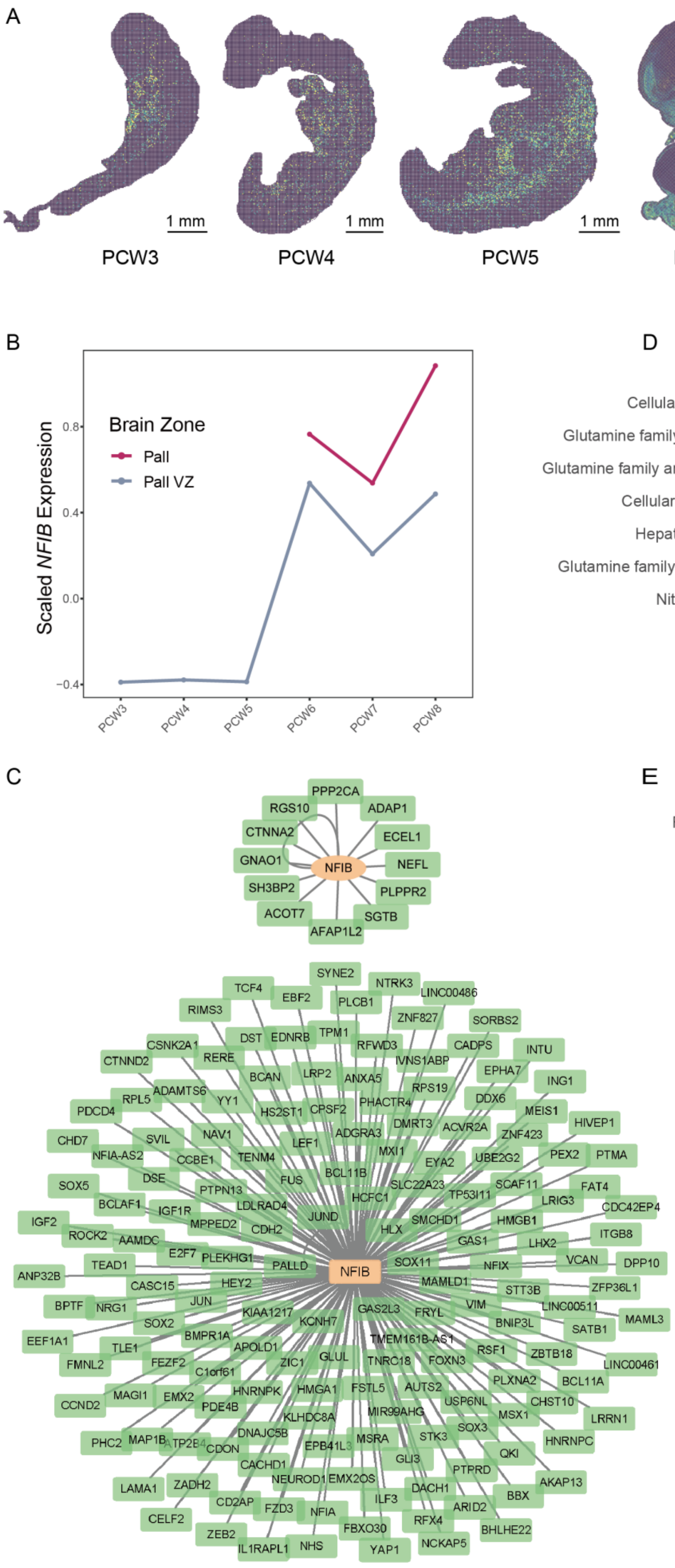
A
1 mm
1 mm
1 mm
2.5 mm
3 mm
5 mm
Max
Min
PCW3
PCW4
PCW5
PCW6
PCW7
PCW8
B
Scaled NFIB Expression
Brain Zone
Pall
Pall VZ
0.8
0.4
0.0
−0.4
PCW3
PCW4
PCW5
PCW6
PCW7
PCW8
D
Arginine metabolic process
Cellular amino acid catabolic process
Glutamine family amino acid catabolic process
Glutamine family amino acid biosynthetic process
Cellular amino acid metabolic process
Hepaticobiliary system development
Glutamine family amino acid metabolic process
Nitrogen cycle metabolic process
Urea metabolic process
Urea cycle
Count
10
20
30
40
50
P.adjust
0.04
0.03
0.02
0.01
PCW5
PCW6
PCW7
PCW8
C
NFIB
PPP2CA
RGS10
ADAP1
CTNNA2
ECEL1
GNAO1
NEFL
SH3BP2
PLPPR2
ACOT7
SGTB
AFAP1L2
E
Collagen trimer
Regulation of vasculature development
Regulation of angiogenesis
Venous blood vessel development
Respiratory system development
Respiratory tube development
Lung development
Positive regulation of endothelial cell migration
Cellular response to VEGF stimulus
VEGF signaling pathway
Regulation of cellular response to growth factor stimulus
Endothelial cell migration
Tissue migration
Ameboidal−type cell migration
Count
10
20
30
40
50
60
P.adjust
0.03
0.02
0.01
PCW8

**Fig. S13. Organ susceptibility to developmental disorders.** (A) Spatial visualization of the expression pattern of *NFIB* from PCW3 to PCW8. (B) Line graph showing *NFIB* expression in Pall and Pall VZ from PCW3 to PCW8. (C) Gene regulatory networks of NFIB in brain at PCW5 (upper) and PCW6 (lower) as visualized by Cytoscape. All targeting genes are shown. (D) Bubbleplot showing the GO enrichment pathways of up-regulated genes in human liver compared with mouse liver from PCW5 to PCW8. (E) Bubbleplot showing the GO enrichment pathways of up-regulated genes in human lung compared with mouse lung at PCW8.

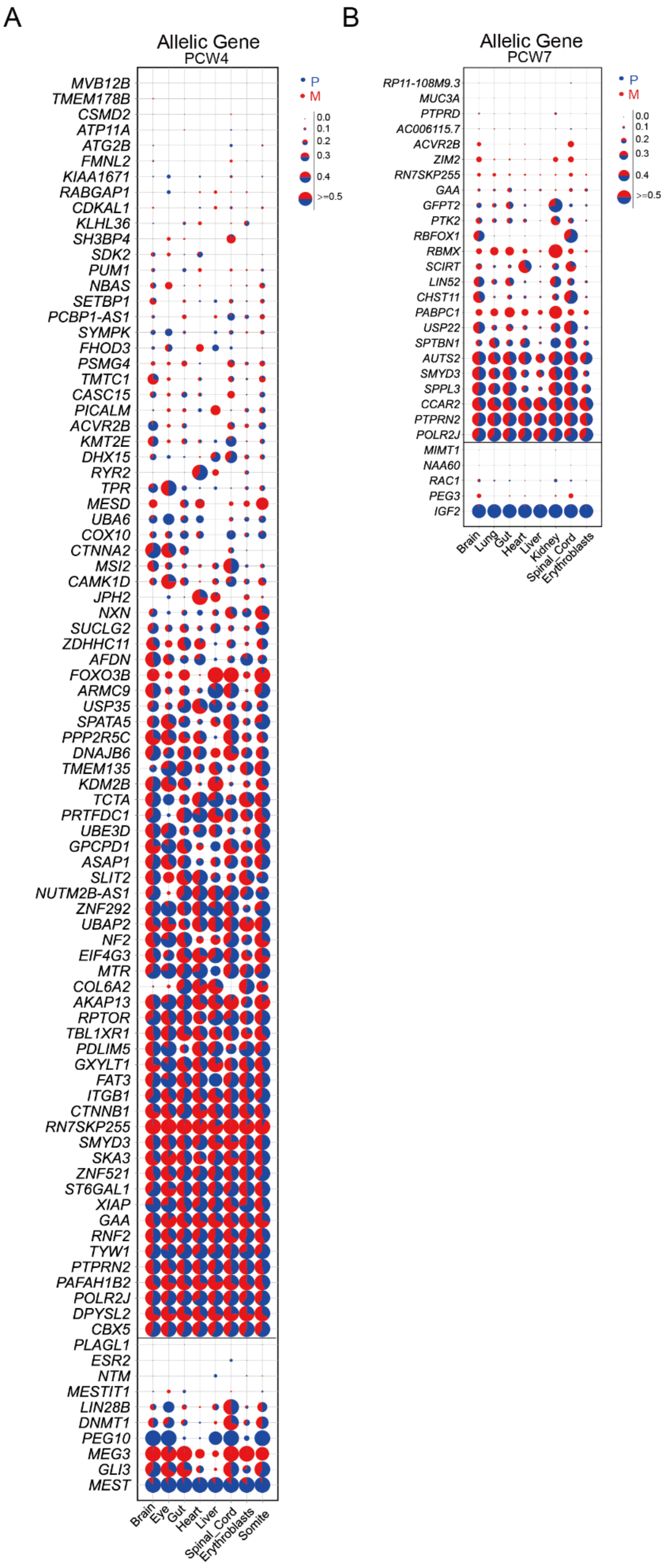
A
Allelic Gene
PCW4
P
M
0.0
0.1
0.2
0.3
0.4
>=0.5
MVB12B
TMEM178B
CSMD2
ATP11A
ATG2B
FMNL2
KIAA1671
RABGAP1
CDKAL1
KLHL36
SH3BP4
SDK2
PUM1
NBAS
SETBP1
PCBP1-AS1
SYMPK
FHOD3
PSMG4
TMTC1
CASC15
PICALM
ACVR2B
KMT2E
DHX15
RYR2
TPR
MESD
UBA6
COX10
CTNNA2
MSI2
CAMK1D
JPH2
NXN
SUCLG2
ZDHHC11
AFDN
FOXO3B
ARMC9
USP35
SPATA5
PPP2R5C
DNAJB6
TMEM135
KDM2B
TCTA
PRTFDC1
UBE3D
GPCPD1
ASAP1
SLIT2
NUTM2B-AS1
ZNF292
UBAP2
NF2
EIF4G3
MTR
COL6A2
AKAP13
RPTOR
TBL1XR1
PDLIM5
GXYLT1
FAT3
ITGB1
CTNNB1
RN7SKP255
SMYD3
SKA3
ZNF521
ST6GAL1
XIAP
GAA
RNF2
TYW1
PTPRN2
PAFAH1B2
POLR2J
DPYSL2
CBX5
PLAGL1
ESR2
NTM
MESTIT1
LIN28B
DNMT1
PEG10
MEG3
GLI3
MEST
Brain
Eye
Gut
Heart
Liver
Spinal_Cord
Erythroblasts
Somite
B
Allelic Gene
PCW7
P
M
0.0
0.1
0.2
0.3
0.4
>=0.5
RP11-108M9.3
MUC3A
PTPRD
AC006115.7
ACVR2B
ZIM2
RN7SKP255
GAA
GFPT2
PTK2
RBFOX1
RBMX
SCIRT
LIN52
CHST11
PABPC1
USP22
SPTBN1
AUTS2
SMYD3
SPPL3
CCAR2
PTPRN2
POLR2J
MIMT1
NAA60
RAC1
PEG3
IGF2
Brain
Lung
Gut
Heart
Liver
Kidney
Spinal_Cord
Erythroblasts

**Fig. S14. The parent-of-origin expressions of 114 phased genes in 8 tissues of PCW4 E3 and PCW7 E3 embryos.**

**Captions Tables for S1 to S21**

**Table S1.**
Summary of Spatiotemporal transcriptomic data.

**Table S2.**
Regulon modules in main fetal organs from PCW3-8.

**Table S3**.
Top 10 regulons in specific organ from PCW3-8.

**Table S4**.
Genes differentially expressed between atrium and ventricle.

**Table S5**.
Gene list of heart sinoatrial node modules.

**Table S6**.
Neural cell markers.

**Table S7.**
Target genes of HMGA2.

**Table S8.**
Enriched expressions of receptors for 14 viruses in specific tissue from PCW3-8.

**Table S9.**
Expression landscape of genes associating with developmental diseases DDG2P in specific tissue from PCW3-8.

**Table S10**.
Enriched genes in certain tissue at specific developmental stages.

**Table S11.**
Target genes of NFIB in brain at PCW5-6, related to figure S13C.

**Table S12**.
Target genes of NFIB at PCW5.

**Table S13**.
GO pathway analysis of NFIB target genes at PCW6.

**Table S14**.
DEGs between embryos of human and mouse in main fetal organs from PCW3-8.

**Table S15**.

GO pathway analysis of up-regulated genes in human liver and lung compared with mouse.

**Table S16**.
The chip on allelic expression analysis across different tissues including, brain, gut, heart, eye, inner ear, liver, lung, kidney and spinal cord.

**Table S17**.
The statistic of overlapped monoallelic expression genes between PCW4 E3 and PCW7 E3.

**Table S18**.
The expressed and distinguishable confidential gene in PCW4 E3 and PCW7 E3.

**Table S19**.
The parent-of-origin differential expression genes in PCW4 E3 and PCW7 E3 and their performance in the other 4 embryos.

**Table S20**.
Sequence information for embryos used for allelic expression analysis.

**Table S21**.
The list of sgRNA sequence for zebrafish cardiac gene knockdown experiments.

**Table S22**.
The list of qPCR primers and probes sequence for zebrafish cardiac gene knockdown experiments.

**Movie S1-S7.**

Pacemaker cell number labeled by Islet-1 (Isl1) in 2 days post fertilization (dpf) zebrafish embryos of control, shox2, roraa, nr2f2, rorab, elapor2b and vsnl1b knockdown group.

**Movie S8-S14.**

Heart rate of zebrafish in control, shox2, roraa, nr2f2, rorab, elapor2b and vsnl1b knockdown group.
